# Quantitative genetics of temperature performance curves of *Neurospora crassa*

**DOI:** 10.1101/2020.01.16.909093

**Authors:** Neda N. Moghadam, Karendeep Sidhu, Pauliina A. M. Summanen, Tarmo Ketola, Ilkka Kronholm

## Abstract

Earth’s temperature is increasing due to anthropogenic CO_2_ emissions; and organisms need either to adapt to higher temperatures, migrate into colder areas, or face extinction. Temperature affects nearly all aspects of an organism’s physiology via its influence on metabolic rate and protein structure, therefore genetic adaptation to increased temperature may be much harder to achieve compared to other abiotic stresses. There is still much to be learned about the evolutionary potential for adaptation to higher temperatures, therefore we studied the quantitative genetics of growth rates in different temperatures that make up the thermal performance curve of the fungal model system *Neurospora crassa*. We studied the amount of genetic variation for thermal performance curves and examined possible genetic constraints by estimating the **G**-matrix. We observed a substantial amount of genetic variation for growth in different temperatures, and most genetic variation was for performance curve elevation. Contrary to common theoretical assumptions, we did not find strong evidence for genetic trade-offs for growth between hotter and colder temperatures. We also simulated short term evolution of thermal performance curves of *N. crassa*, and suggest that they can have versatile responses to selection.

## Introduction

Earth’s temperature is rising due to anthropogenic activities (IPCC, 2013). The challenge most organisms will face in a warming world is that they have to either adapt to warmer conditions or migrate into colder areas to avoid extinction (Deutsch et al., 2008; Dillon et al., 2010; Araújo et al., 2013; Merilä and Hendry, 2014). Temperature is a unique abiotic stress, because the kinetics of all biochemical reactions and protein stability are affected by temperature. As such, temperature influences nearly all aspects of an ectothermic organism’s physiology (Schulte, 2015; Arcus et al., 2016). Therefore, adapting to a higher temperature may be much more difficult than adapting to a more specific environmental stress. For some anthropogenic stresses, such as antibiotics or herbicides, decades of research have revealed strong evolutionary adaptation to these stresses (Davies and Davies, 2010; Powles and Yu, 2010). However, genetic basis of adaptation to temperature is likely to be much more complex (Hochachka and Somero, 2002).

According to quantitative genetic theory, evolution is possible if variation in a trait is heritable and selection acts on this variation. However, the evolution of multivariate traits can be complicated by genetic correlations, allowing evolution to proceed only in few directions or possibly preventing it altogether (Walsh and Blows, 2009). The more integrated traits are with each other, the more difficult the evolution of the underlying genetic network and the phenotype can be.

The ability of an organism to tolerate different temperatures is often described by a thermal performance curve (Huey and Kingsolver, 1989, 1993), which describes the fitness or performance of an organism as a function of temperature (Figure 1A). These curves have been used to predict how organisms potentially respond to increased temperatures (Deutsch et al., 2008; Araújo et al., 2013; Sinclair et al., 2016). In general, thermal performance curves or reaction norms have been thought to evolve by either changes in elevation (Figure 1B), left or right shifts in the curve that lead to changes in optimum temperature or temperature limits (Figure 1C), or changes in curve shape (Figure 1D).

**Figure 1:**
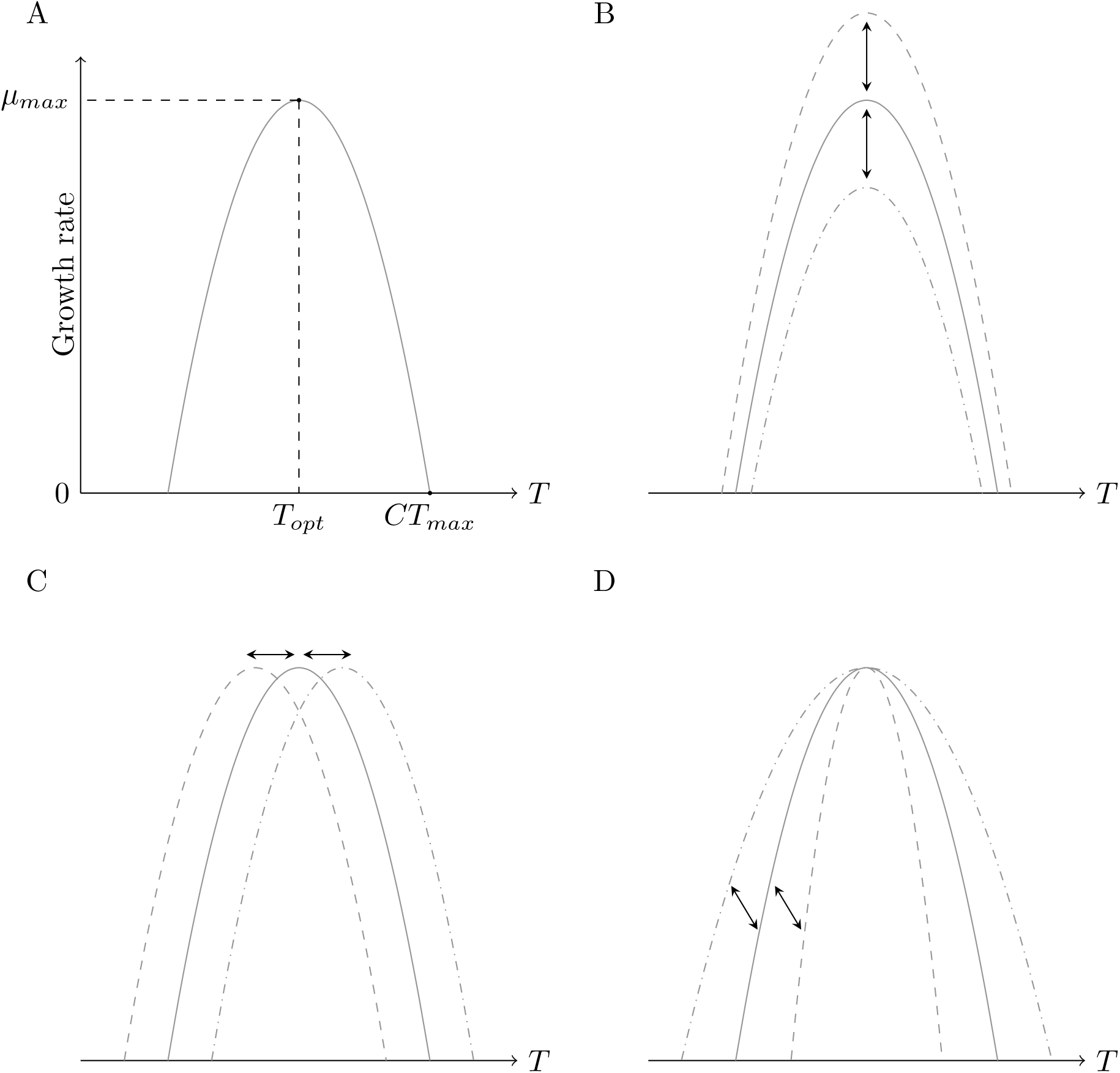
A) An illustration of a hypothetical temperature performance curve. Temperature is on the horizontal axis and growth rate is on the vertical axis. *T_opt_* shows the optimal temperature, where growth rate has its maximum value, *μ_max_*. Temperature where growth rate reaches zero as temperature increases is denoted as *CT_max_*. B) Change in reaction norm elevation shifts the reaction norm on the vertical axis. C) A horizontal shift. D) Change in reaction norm shape.

Certain biochemical constraints may explain the characteristic shape changes of performance curves (Angilletta et al., 2003). For example, high enzyme stability could allow tolerating high temperatures with the expense of reduction of performance in cold temperatures resulting in a hot-cold trade-off. Two enzymes with different optima could allow broader thermal tolerance but with an energetic expense of expressing two proteins, leading to reduction of performance at intermediate temperatures and producing a specialist-generalist tradeoff. Furthermore, the biochemial activation energy provided by higher temperatures can lead to thermodynamic effects: genotypes with higher optimal temperatures also have higher performance (Hochachka and Somero, 2002). Thermodynamic effect is also called the “hotter is better” hypothesis. If thermal performance curves are determined by such underlying patterns, measurements need to be done in multiple temperatures and results analyzed by multivariate methods in order to determine the ability of thermal performance to evolve.

While several studies have tested how different species or populations differ in their thermal performance curves, or if evolution has been able to shape them (e.g. Krenek et al., 2011; Klepsatel et al., 2013; Ketola and Saarinen, 2015; Ashrafi et al., 2018; Maclean et al., 2019), only a few studies have determined the evolutionary potential of thermal performance curves. In these studies, the genetic variance-covariance matrix (**G**-matrix) for thermal performance across several temperatures has been estimated, and how genetic variation is aligned with characteristic directions of reaction norm evolution has been determined (e.g. Izem and Kingsolver, 2005; Stinchcombe et al., 2010; Latimer et al., 2015; Logan et al., 2020). This is essential in order to explore how freely thermal performance can evolve in different environments, and to quantify if thermal performance evolution is bound to follow a certain evolutionary path or performance curve shape. Constraints on performance curve evolution will affect the ability of populations to respond to increasing temperatures, which is crucial, as studies suggest that plastic responses alone may not be enough for most species for dealing with coming temperature increases (Gunderson and Stillman, 2015).

However, in the midst of multivariate genetics and the emphasis on finding genetic constraints, it should be remembered that evolutionary change follows from selection. From quantitative genetic parameters one can only deduce which traits have the highest amount of variation, and what is the alignment of the **G**-matrix with respect to characteristic thermal performance curve shapes. However, unless genetic correlations are exactly −1 or 1, or if selection occurs exactly to the direction of zero genetic variation, evolutionary change to a particular direction is not prohibited, only slowed down.

To explore constraints of thermal performance curve evolution, we are using the filamentous fungus *Neurospora crassa* as a model system to study the quantitative genetics of thermal performance curves. *N. crassa* is a genetic model system that has been used extensively in different aspects of genetic research (Roche et al., 2014). However, only recently some studies have started to explore quantitative variation in *N. crassa* (Ellison et al., 2011; Palma-Guerrero et al., 2013). This is despite *N. crassa* having excellent properties for the study of quantitative genetics: *N. crassa* can reproduce either asexually or sexually, so analysis of clones is possible for quantitative genetic experiments and controlled crosses can be made. Comparatively little is known about the ecology of *N. crassa*; it is a saprotrophic organism that decomposes dead plant matter, and it is particularly found on burned vegetation. Its geographic distribution is concentrated in mainly tropical and subtropical regions (Turner et al., 2001). Most strains have been collected from the Caribbean basin, southeastern United States, west Africa, and India (Turner et al., 2001), but the species also occurs in southern Europe (Jacobson et al., 2006).

Specifically, we asked the following questions: 1) Is there genetic variation in thermal performance curves in *N. crassa*? 2) Is variation in performance curves mainly for elevation, location, or shape? 3) Do constrains exist for performance curve evolution in the short term and what are these constraints?

To address how much there is genetic variation in temperature performance curves, we used a panel of strains of *N. crassa* that had earlier been sampled from natural populations. We also crossed certain strains together to generate additional families. We measured the growth rates of these strains in different temperatures and combine these measurements into a thermal performance curve, and used a multivariate model to estimate the **G**-matrix of performance at different temperatures. We then used the empirical estimates of genetic variation in a quantitative genetic model to describe the short term evolutionary potential of temperature performance curves of *N. crassa*.

## Materials and methods

### *Neurospora crassa* strains

We used a panel of strains originally obtained from the Fungal Genetics Stock Center (McCluskey et al., 2010). Our sample included natural strains collected from Louisiana (USA), Caribbean, and Central America (Ellison et al., 2011; Palma-Guerrero et al., 2013), 113 natural strains in total. In addition we made crosses between some of the strains to obtain additional families and increase the amount of genetic variation segregating among our lines. We crossed strains 10948 × 10886 to obtain family A (*n* = 94), 10932 × 1165 to obtain family B, (*n* = 50), 4498 × 8816 to obtain family C (*n* = 50), 3223 × 8845 to obtain family D, (*n* = 52), and 10904 × 851 to obtain family G (*n* = 69). Parents were chosen to have crosses within the Louisiana strains and between the Louisiana and Caribbean strains. In total, the panel contained 428 strains and based on genotypic data (Ellison et al., 2011; Palma-Guerrero et al., 2013) all strains represent unique genotypes. Table S1 contains a list and information about the used strains. Strain numbering in family G runs up to 72, because strains G2, G9, and G51 grew very poorly and were excluded from the experiment.

### Phenotyping

Standard laboratory methods were used to maintain *Neurospora* cultures (Davis and de Serres, 1970). We measured growth rates using a tube method described in Kronholm et al. (2016) but instead of parafilm we used silicone plugs to cap the tubes. We measured the linear growth rate of each genotype in six different temperatures: 20, 25, 30, 35, 37.5, and 40 °C. Temperatures were chosen based on known reaction norm for strain 2489 (Kronholm et al., 2016). Three clonal replicates were measured for each genotype at each temperature. This gave a total of 7704 growth assays. In some assays the inoculation failed and the strain did not grow, or water droplets moved the inoculum along the pipette and linear growth rate could no longer be measured. There were 19 such assays and these were recorded as missing data, thus the number of growth assays in the final dataset was 7685. Strains were grown in two growth chambers (MTM-313 Plant Growth Chamber, HiPoint Corp., Taiwan) that contained three compartments, each with adjustable temperature. We rotated the temperatures among the different compartments between replicates, so that replicates of the same temperature were measured in different compartments, and monitored the temperature in the compartments with data loggers.

### Statistical analysis

All statistical analyses were performed with R 3.6.0 (R Core Team, 2019). Bayesian models were implemented using the Stan language (Carpenter et al., 2017) which uses Hamiltonian Monte Carlo sampling. Hamiltonian Monte Carlo is much more efficient than traditional MCMC algorithms, such as Gibbs sampling, and can potentially accommodate very large number of parameters. An accessible introduction can be found in McElreath (2015). Stan was interfaced using the ‘brms’ 2.9.0 R package (Bürkner, 2018). MCMC convergence was monitored by trace plots and 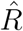 values. We considered parameter values to be different if their 95% highest posterior density (HPD) intervals did not overlap.

#### Thermodynamics of thermal performance curves

Theory predicts that if differences between hot and cold adapted genotypes are determined solely by an effect of temperature on metabolic rate, named the thermodynamic effect or “hotter is better” hypothesis, there should be a negative relationship between the logarithm of maximal growth rate, *μ_max_*, and 1/(*kT_opt_*), where *k* is the Boltzmann’s constant, and *T_opt_* is the temperature (in K) at which maximal growth rate occurs (Savage et al., 2004). We examined whether differences between *N. crassa* genotypes could be solely explained by a thermodynamic effect. When ln(*μ_max_*) is plotted against 1/(*kT_opt_*) the slope of a regression line is equal to negative activation energy, −*E*. The thermodynamic expectation for the slope is −0.6 because 0.6 eV is the average activation energy for biochemical reactions in the cell. This pattern generally holds across taxa adapted to different temperatures (Savage et al., 2004; Sørensen et al., 2018). Slopes greater than −0.6 have been interpreted as an indication of other physiological or biochemical reasons rather than a thermodynamic effect (Sørensen et al., 2018).

To calculate the optimum temperature for each genotype without using a specific model that may fit for some genotypes better than others, we fitted natural splines for each genotype. We extracted the maximum growth rate (*μ_max_*) and optimum temperature (*T_opt_*) from the spline fit. We then fit a model

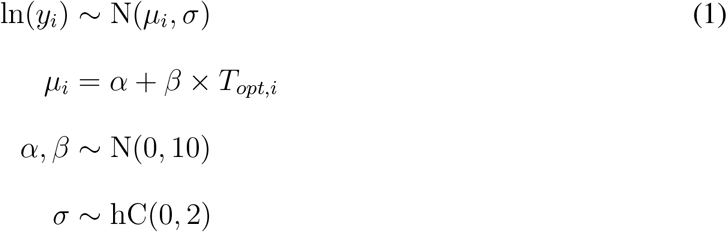

where *y_i_* is the *i*th maximum growth rate, *α* is the intercept, *β* is the slope, and *T_opt,i_* is the *i*th optimum temperature. We used weakly regularizing priors: a normal distribution for *α* and *β*, and a half-cauchy distribution for *σ* with location 0 and scale 2. MCMC estimation was done using two chains, with a warmup of 1000 iterations, followed by 4000 iterations of sampling. For this analysis we removed genotypes from the data that had very low maximal growth rates ln(*μ_max_*) < 1, which is *μ_max_* < 2.72 mm/h, as they did not have the typical tolerance curve shape and were outliers. These genotypes typically grew very slowly and reaction norms were much flatter t han typical ones, which leads to larger uncertainty in estimating the optimal temperature from the spline fits (Figure S1A). 14 genotypes were removed, this left 414 genotypes for the analysis. However, since removing outlier observations can be considered subjective, we also applied robust regression with bisquare weights to the full data. Robust regression is a method that gives less weight to individual data points than ordinary regression and is less affected by outlier observations (Venables and Ripley, 2002).

#### Estimation of genetic variance and covariance components

We were interested in estimating the genetic variance and covariance components for growth rates at different temperatures that together describe different aspects of temperature performance curves. Because *Neurospora* can be propagated clonally, we can estimate genetic variance components using clonal analysis. Among genotype variance is an estimate of the genetic variance and within genotype variance is an estimate of the environmental variance (Lynch and Walsh, 1998). We used a multivariate model to estimate genetic variance components at each temperature and the genetic correlations of all possible temperature pairs. The advantage of this approach is that we do not have to assume any particular shape for the temperature reaction norm. The multivariate model was

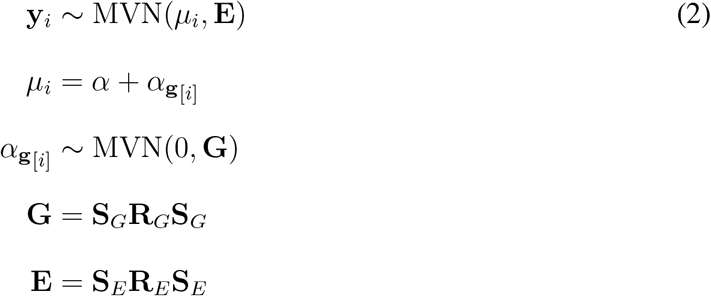

where *α* is a vector of intercepts, 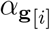 is a vector of genotypic effects, **S**_*G*_ and **S**_*E*_ are 6 × 6 diagonal matrices with genetic or environmental standard deviations on the diagonal, and **R**_*G*_ and **R**_*E*_ are matrices for genetic and environmental correlations respectively. We used weakly informative priors by using the half location-scale version of the student’s t distribution with three degrees of freedom and 10 as the scale parameter. Thus, the prior for intercept effects was

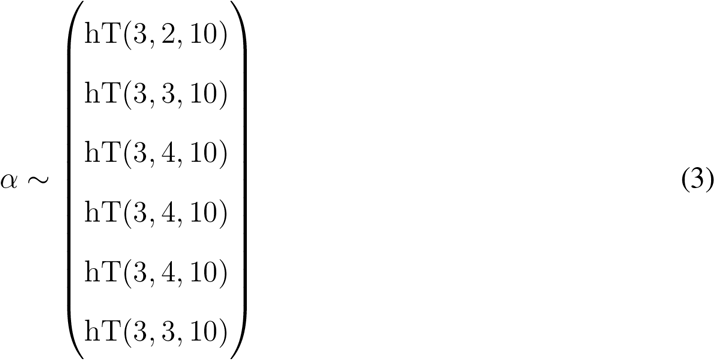

for growth rates from 20 to 40 °C. The prior for each standard deviation in the model was *σ* ∼ hT(3, 0, 10), and we used an lkj prior for the correlation matrices: **R**_*E*_, **R**_*G*_ ∼ LKJ(1). For MCMC estimation two chains were run with a warmup period of 1000 iterations, followed by 5000 iterations of sampling, with thinning set to 2. By inspecting MCMC traceplots (Figure S2) and the diagnostic summary statistic 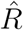, which was 1 for all parameters, we found no evidence of convergence problems.

#### Genetic correlations and temperature differences

We were also interested in how the genetic correlation of growth rates changes as temperatures are further apart. In order to examine how correlations change in a statistically rigorous manner, we calculated pairwise temperature differences for each estimated genetic correlation (*n* = 15), and fitted a Bayesian linear model with genetic correlation as the response, taking into account the uncertainty in the estimated genetic correlations. This is a linear model with measurement error where uncertainty in the estimated genetic correlations is propagated to the intercept and slope estimates of the linear model, see McElreath (2015) for details. We compared models with or without slope effects for temperature and whether genetic correlations involving growth rate at 40 °C had a different intercept or slope (Table 2). We used leave-one-out cross-validation method for model comparisons, implemented in the ‘loo’ R package (Vehtari et al., 2017). The models were compared using the leave-one-out information criterion; smaller values indicate greater support for a model. The final model was

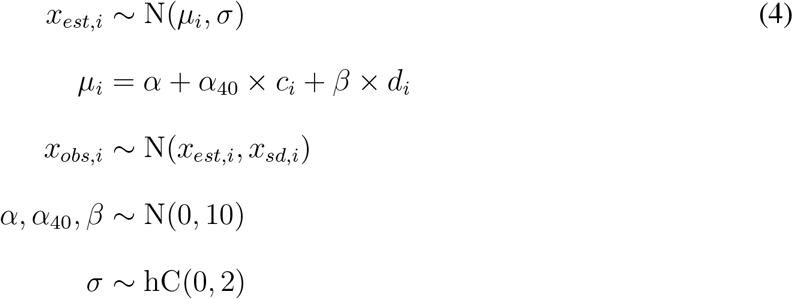

where *x_obs,i_* is the median of *i*th observed genetic correlation, *x_sd,i_* is the observed standard deviation of *i*th genetic correlation, *x_est,i_* is the estimated genetic correlation for *i*th observation, *α* is the intercept, *α*_40_ is the intercept effect when one of the temperatures is 40 °C, *c_i_* is an indicator variable whether one of the temperatures is 40 °C, *β* is the slope effect, and *d_i_* is the temperature difference for the *i*th observation. MCMC estimation was done using two chains, with a warmup of 1000 iterations, followed by 4000 iterations of sampling.

#### Quantitative genetics

We estimated heritability, the proportion of genetic variance of the total variance, for each temperature as

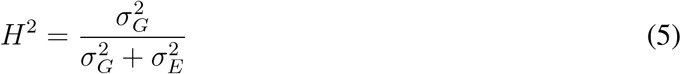

where 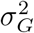 is the genetic variance component and 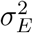 the environmental variance component. Because *Neurospora* is haploid, the dominance variance component is not defined. Genetic variance in haploids is composed of

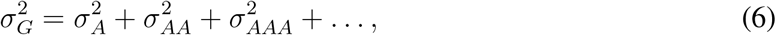

where 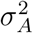 is the additive variance and 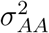 is the additive × additive epistatic variance, 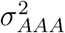 is the additive × additive × additive variance, and so on (Lynch and Walsh, 1998). With our experimental design we cannot estimate the epistatic variance terms, as is the case with many other common quantitative genetic designs, and going further we assumed that epistatic variances were small and were ignored. This seems like a strong assumption, but there is some justification for doing so: even if there is plenty of epistasis at the level of gene action, this is not necessarily translated into epistatic variance (Hill et al., 2008; Mäki-Tanila and Hill, 2014). Empirical data also suggest that most genetic variation is additive (Hill et al., 2008). The genetic covariance of traits 1 and 2 is 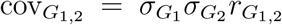, where 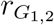 is the correlation of the standard deviations or the genetic correlation. Thus, genetic correlation for traits 1 and 2 can be defined as

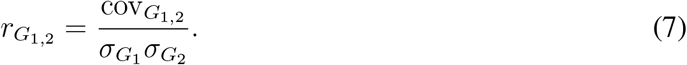

In addition to heritabilities, we used coefficients of variation to compare genetic and environmental variances. Heritability can be influenced by changes in either genetic or environmental variance, and genetic variance by itself is not a unitless variable (Houle, 1992). The genetic coefficient of variation was:

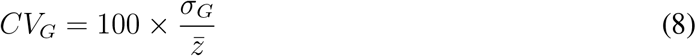

where 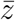 is the mean phenotype. Accordingly, the environmental coefficient of variation was 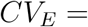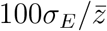.

We obtained the **G**-matrix to describe how the growth rates at different temperatures were correlated and to be able to calculate multivariate response to selection for thermal performance curves. This matrix contains genetic variance components on the diagonal and covariance components on off-diagonals, so for *n* traits **G** is an *n* × *n* matrix:

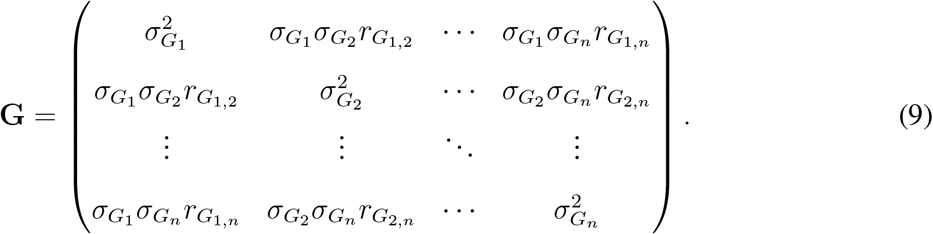

For environmental variance it is possible to construct an analogous **E**-matrix that is the environmental variance-covariance matrix.

We performed eigen decomposition of the **G**-matrix to gain insight into genetic constraints of reaction norm evolution. The eigenvector corresponding the leading eigenvalue, or the first principle component, gives the direction of multivariate evolution with the least genetic resistance (Schluter, 1996). We obtained these components by principle component analysis of the **G**-matrix. To assess uncertainty in the eigen decomposition we constructed a **G**-matrix for each posterior sample and performed decomposition for each **G**-matrix to obtain posterior distributions for how much variance the different components explained and for the component loadings. Obtaining interval estimates for the loadings this way is valid only if the order of eigenvectors stays consistent between the samples, and we could confirm this for the components one and two.

To assess evolvability and constraint across the different growth rates we used the approach of Hansen and Houle (2008). Assuming there is a directional selection gradient ***β*** in multivariate space, they define evolvability as the length of the response to selection in the direction of ***β***, this is the same as projection of response to selection on ***β*** (Hansen and Houle, 2008). Evolvability was calculated as

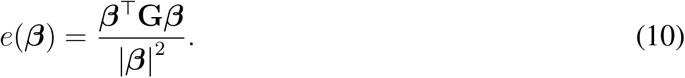

Furthermore they define conditional evolvability as the response to selection in the direction of ***β***, assuming that there is stabilizing selection around the direction of ***β*** and the population cannot deviate from this direction. For conditional evolvability we first calculated unit vector of ***β*** as

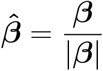

and conditional evolvability is then

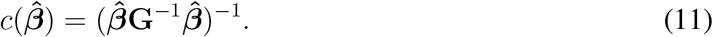

To asses whether evolvability along a certain selection gradient is particularly high or low it is possible to calculate average evolvabilities over random selection gradients in phenotypic space. Hansen and Houle (2008) derived analytical and approximate solutions for average evolvability and average conditional evolvability and we calculated these following their approach. Evolvabilities for single traits are just the genetic variances of those traits. Conditional evolvability for a single trait can be measured with respect to other traits. Conditional evolvability for trait *i* is *c_i_* = 1/[**G**^−1^]_*ii*_, where [**G**]_*ii*_ is the *i*th diagonal element of the **G**-matrix. Trait autonomy, the proportion of evolvability that remains after conditioning for the other traits, is calculated as *a_i_* = ([**G**^−1^]_*ii*_[**G**]_*ii*_)^−1^ (Hansen and Houle, 2008). Since there are scale differences in the growth rates at different temperatures, we calculated conditional evolvabilities for both on the original scale and on mean standardized scale. The **G**-matrix can be mean standardized by dividing *ij*th element by the product of the means of traits *i* and *j*. 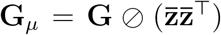, where 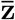 is a vector of trait means and ⊘ symbol for elementwise division. The mean standardized selection gradient was calculated as 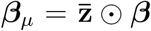, where ⊙ is element-wise multiplication. Interval estimates for these statistics were obtained by calculating them for each posterior sample.

#### Quantitative genetic model for the evolution of performance curves

To examine how thermal performance curves of *N. crassa* can evolve, we used a quantitative genetic model with the empirically estimated **G**-matrix. Response to selection can be calculated using the multivariate breeder’s equation

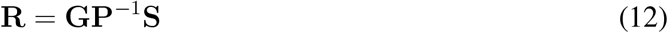

where **S** is a vector of selection differentials for each temperature, **G** and **P** are the genetic and phenotypic variance-covariance matrices respectively, and **R** is the response to selection. Response to selection can also be expressed in terms of the selection gradient, ***β***, as

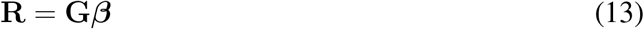

where ***β*** = **P**^−1^**S**. The biological interpretation of selection differential and selection gradient are different, as selection differential of 0 for a given trait does not imply selective neutrality but rather stabilizing selection, whereas selection gradient of 0 for a trait implies that the trait is selectively neutral. See figure S3 for an illustration of the differences between these concepts. When we asked how would evolution proceed in a particular direction, we simulated response to selection using selection gradient (Equation 13). And when we asked whether selection could generate a particular phenotype we simulated response to selection using selection differentials (Equation 12). Our goal is not to predict the evolution of tolerance curves in nature, as the real selection gradients are unknown and the assumption that **G** remains constant is likely violated in real populations. Indeed, there are considerable difficulties in predicting the response to selection in nature (Morrissey et al., 2010). Instead, our goal is to illustrate how thermal performance curves could evolve in a population with a similar **G** as estimated empirically here.

The phenotypic matrix was calculated from **P** = **G**+**E**. The environmental variance-covariance matrix **E**, which uses environmental standard deviations and their correlations analogous to equation 9, was obtained from the same model fit as **G**. Since there is uncertainty in our estimates of **G** and **E** we incorporated this uncertainty in the selection responses by sampling 1000 **G** and **E** matrices from the posterior distributions of genetic and environmental standard deviations and calculating a response to selection for each sample. We calculated responses to selection after 1, 3, and 5 generations of selection, assuming that the selection differentials, **G**, and **E** matrices stay the same. We always normalized the sum of absolute values of selection differentials or gradients across all temperatures for different selection regimes. First we used selection gradients that corresponded to the first two eigenvectors of the **G**-matrix. The summed absolute values of selection gradients or selection differentials across all temperatures were normalized to be 0.6 mm/h. We estimated evolvability and conditional evolvability along these gradients as explained above. Then we used different selection regimes to examine how we could change the performance curve elevation, optimum, or shape (Figure 1). We used six different vectors of **S**: **S**_1_ = {0.1, 0.1, 0.1, 0.1, 0.1, 0.1} and **S**_2_ = {−0.1, −0.1, −0.1, −0.1, −0.1, −0.1}, which correspond to selection on elevation change; **S**_3_ = {0, 0.1, 0.1, −0.2, −0.2, 0} and **S**_4_ = {0, −0.05, −0.05, −0.1, 0.2, 0.2}, which correspond to a shift in optimum temperature; **S**_5_ = {0.1, 0.2, 0, 0, 0.1, 0.2} and **S**_6_ = {0, −0.1, −0.25, 0.05, −0.1, −0.1}, which correspond to change in reaction norm shape. The selection differentials were chosen so that they would produce the desired phenotypic change, choice of numerical values was otherwise arbitrary. For evolvability calculations, we calculated realized selection gradients based on these selection differentials as ***β*** = **P**^−1^**S**.

## Results

### Growth of *Neurospora* at different temperatures

Temperature had a large effect on growth, at 20 °C growth rate was between 2 and 2.5 mm/h (mean 2.17 and 95% HPD interval 2.15–2.20) for most strains, and as temperature increased up to 35 °C growth rates rose to between 3 and 5 mm/h (mean 4.15 and 95% HPD interval 4.08–4.22) for most strains (Figure 2A). This represents an increase of 91% in mean growth rate. For many strains growth rate peaked at 35 °C and then decreased as temperature was increased (Figure 2A), at 40 °C mean growth rate was 2.35 (2.29–2.41 95% HPDI) mm/h. The performance curves of *N. crassa* exhibited a typical performance curve form: with an optimum temperature and decrease in growth rate in other temperatures, and performance declined faster in temperatures warmer than the optimum (Sinclair et al., 2016). Few genotypes grew very slowly and had unusual tolerance curve shapes (Figure 2A), possibly reflecting that these genotypes were poorly suited to lab conditions, due to specific nutritional requirements for example.

**Figure 2:**
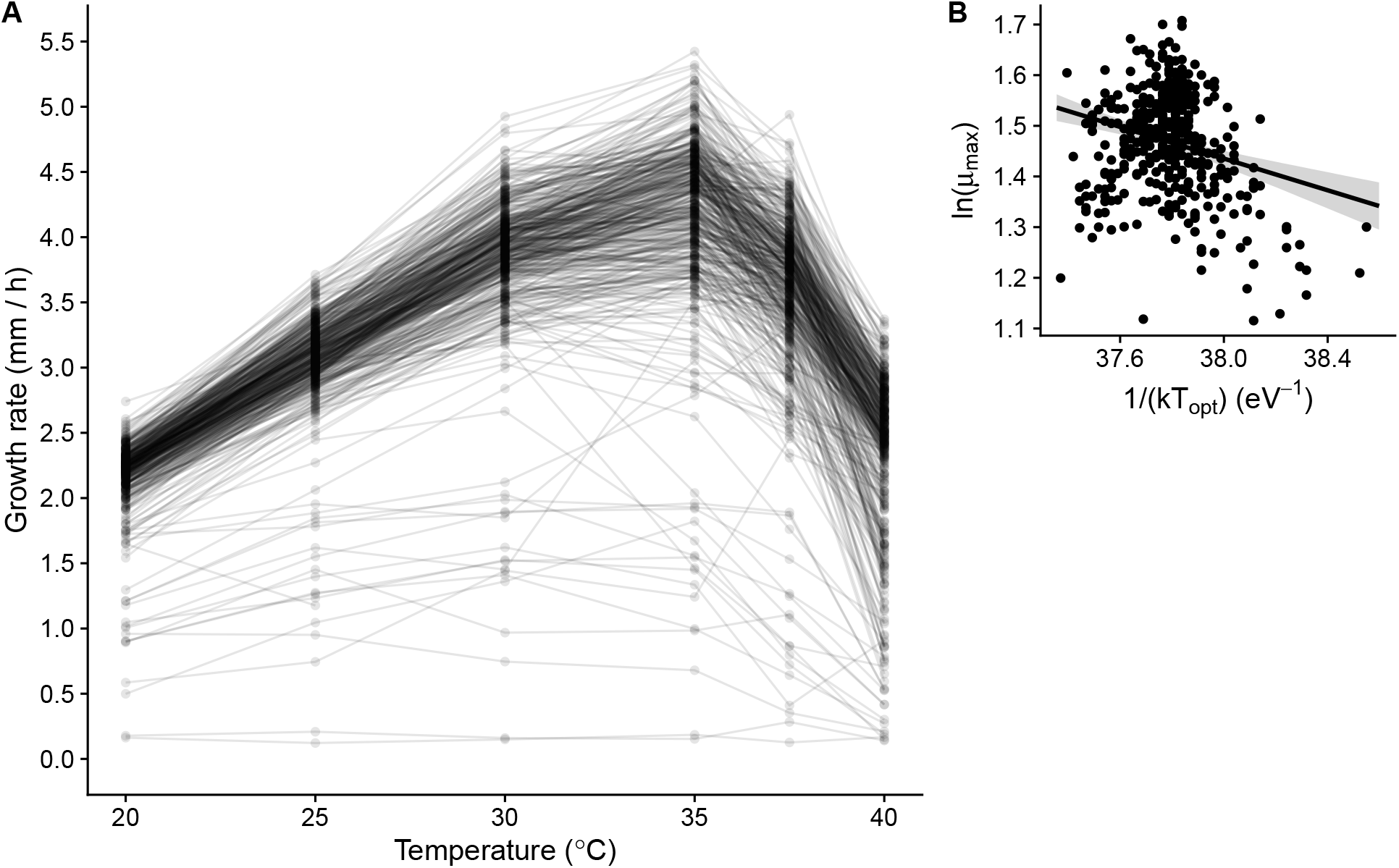
A) Phenotypic means for each genotype. B) Logarithm of maximum growth rate, *μ_max_*, plotted against inverse of *kT_opt_*, where *k* is the Boltzmann’s constant and *T_opt_* the temperature where maximal growth rate occurs. The slope gives an estimate of negative activation energy −*E*.

### Thermodynamics of thermal performance curves

We examined whether differences between genotypes could be explained by a thermodynamic effect, i. e. does the maximum growth rate increase with optimum temperature. We obtained *μ_max_* and *T_opt_* from the natural spline fits and plotted ln(*μ_max_*) against 1/(*kT_opt_*) (Figure 2B). For the bulk of the genotype data, the estimated slope was −0.16 (95% HPD from −0.22 to −0.10), which corresponds to activation energy of 0.16 eV. This was lower than the theoretical expectation of 0.6 eV. Moreover, there was substantial amount of variation around the regression line (Figure 2B); optimum temperature explains only a small proportion of the observed variation. This indicates that while a small thermodynamic effect exists, most variation within *N. crassa* is due to other physiological and biochemical causes. As this result was obtained in an analysis where we removed genotypes which had atypical reaction norms (Figure S1A), we also fitted a robust regression to the full data (Figure S1B) and obtained a slope of −0.17 which is very close to our original estimate of −0.16. While fitting an ordinary regression to the full data gives a somewhat smaller slope (−0.34), the few atypical observations have high leverage in the model. Since results of robust regression and removing outliers agree, it seems that removing the outliers is quite reasonable in this case.

### Quantitative genetics

In order to analyse the data without forcing the tolerance curves to fit any predetermined shape, or underlying latent structures as in Izem and Kingsolver (2005), we fit a multivariate model to the data where growth at each temperature was modelled as potentially correlated with growth at other temperatures. We obtained the **G**-matrix from the multivariate model fit (Equation 2). There was genetic variation for growth in all temperatures and all genetic covariances and correlations were positive (Table 1).

**Table 1:** Genetic variances, covariances, correlations, and environmental variances for growth rates in different temperatures estimated from the multivariate model. The diagonal (in bold) contains genetic variances 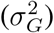, upper triangle contains genetic covariances 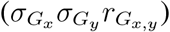, and lower triangle contains genetic correlations 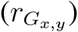. The last column contains environmental variances 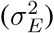. Estimates are posterior means with 95% HPD intervals shown in parenthesis.

| (°C) | 20 | 25 | 30 | 35 | 37.5 | 40 | $\sigma_E^2$ |
| --- | --- | --- | --- | --- | --- | --- | --- |
| 20 | **0.08 (0.07–0.09)** | 0.11 (0.1–0.13) | 0.14 (0.12–0.16) | 0.15 (0.13–0.18) | 0.13 (0.11–0.15) | 0.07 (0.05–0.09) | 0.01 (0.01–0.01) |
| 25 | 0.94 (0.93–0.96) | **0.17 (0.15–0.19)** | 0.22 (0.19–0.25) | 0.24 (0.21–0.28) | 0.19 (0.16–0.22) | 0.11 (0.09–0.14) | 0.02 (0.02–0.02) |
| 30 | 0.86 (0.83–0.89) | 0.96 (0.94–0.97) | **0.32 (0.28–0.36)** | 0.36 (0.31–0.41) | 0.28 (0.23–0.32) | 0.17 (0.13–0.21) | 0.04 (0.03–0.04) |
| 35 | 0.75 (0.7–0.79) | 0.83 (0.79–0.86) | 0.9 (0.88–0.92) | **0.5 (0.43–0.57)** | 0.42 (0.36–0.48) | 0.25 (0.2–0.3) | 0.06 (0.05–0.07) |
| 37.5 | 0.68 (0.62–0.73) | 0.7 (0.65–0.75) | 0.75 (0.7–0.8) | 0.9 (0.88–0.92) | **0.43 (0.37–0.49)** | 0.3 (0.25–0.35) | 0.09 (0.08–0.1) |
| 40 | 0.42 (0.33–0.51) | 0.47 (0.39–0.56) | 0.51 (0.43–0.59) | 0.61 (0.54–0.68) | 0.77 (0.72–0.82) | **0.34 (0.29–0.4)** | 0.2 (0.18–0.22) |

**Table 2:** Comparison of different models for relationship between genetic correlations and temperature differences. Model terms correspond to different deterministic parts of the model in equation 4, *α*_40_ is an intercept effect for correlations involving 40 °C and *β*_40_ is a slope effect for correlations involving 40 °C. LOOIC = Leave-one-out information criterion. SE = standard error.

| Model terms | LOOIC | diff (±SE) | weight |
| --- | --- | --- | --- |
| $\alpha + \alpha_{40} \times c_i + \beta \times d_i$ | -38.54 | 0 (0) | 0.84 |
| $\alpha + \alpha_{40} \times c_i + \beta \times d_i + \beta_{40} \times d_i \times c_i$ | -35.18 | 3.36 (1.24) | 0.16 |
| $\alpha + \beta \times d_i + \beta_{40} \times d_i \times c_i$ | -26.07 | 12.47 (5.49) | 0 |
| $\alpha + \beta \times d_i$ | -14.12 | 24.42 (5.54) | 0 |
| $\alpha$ | -7.61 | 30.93 (6.34) | 0 |

By plotting the model means and genetic correlations it appeared that genetic correlation between adjacent temperatures was generally high, and decreased as temperatures were further apart and correlations involving 40 °C also seemed lower (Figure 3A). We tested this idea formally and fitted a model of genetic correlations and temperature differences. We compared the different models, and the best model had different intercepts for correlations involving 40 °C and for correlations not involving 40 °C, and identical slopes for these two groups (Table 2). A model with both different slopes and different intercepts had marginal weight in the model comparison but the *β*_40_ parameter had an estimate overlapping with zero, so this model gave the same results as the simpler model and thus we report results only from the different intercepts model. The model confirmed our observation that the genetic correlation between any two temperatures was indeed lower if one of those temperatures was 40 °C (Figure 3B), the intercept effect *α*_40_ had an estimate of −0.24 (with a 95% HPDI from −0.31 to −0.17). The genetic correlation of growth rates in two temperatures decreased by 0.02 (0.02–0.01 95% HPDI) units as temperature difference increased by 1 °C. This result suggested that variation in different genes contributes to genetic variation for growth at 40 °C than in lower temperatures.

**Figure 3:**
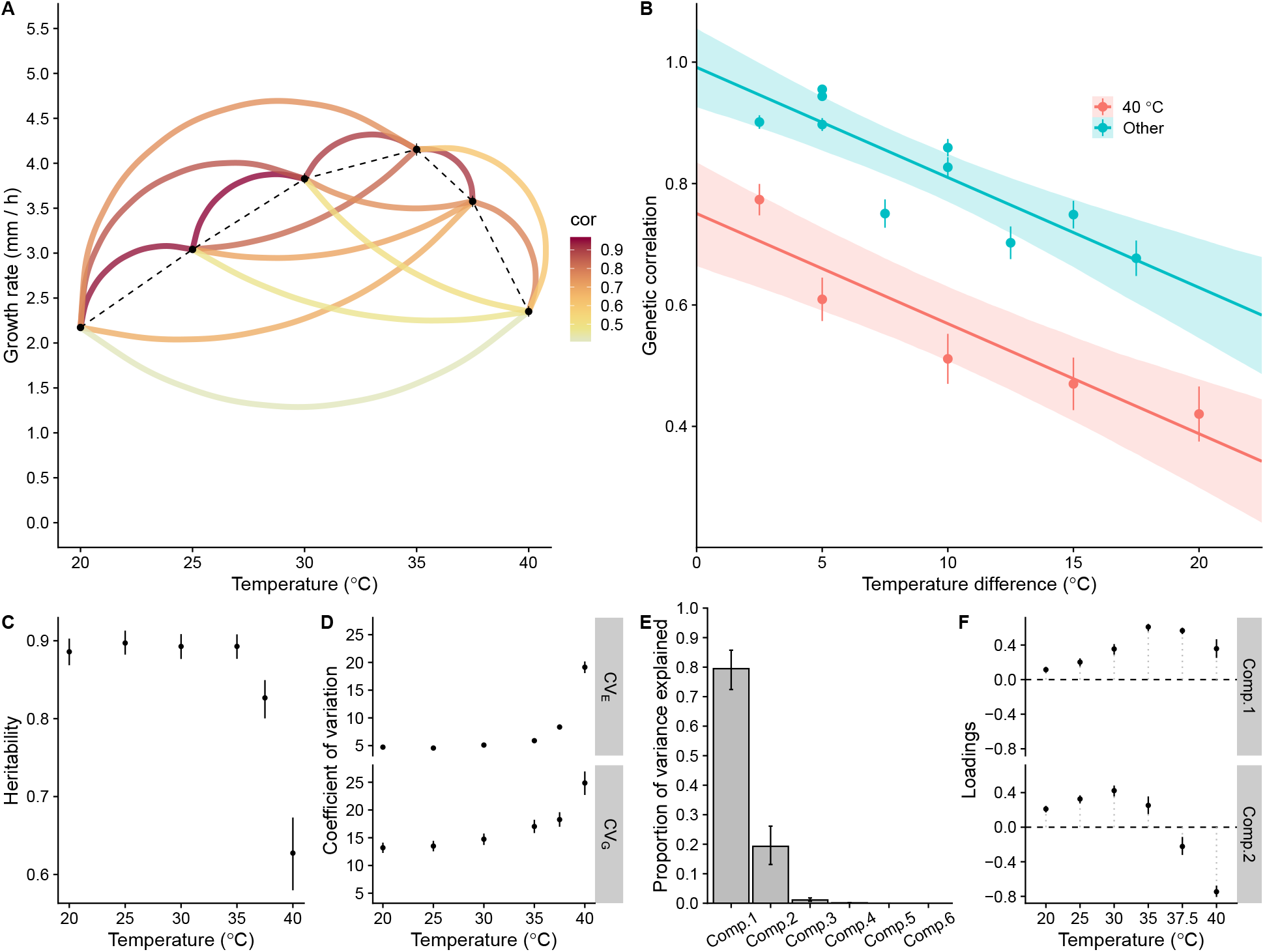
A) Model means and genetic correlations for each temperature. Arcs connect each pair of temperatures and arc color corresponds to the strength of their genetic correlation. B) Genetic correlations against temperature differences. Line is the mean slope of the model and envelope the 95% HPD interval for the slope. C) Heritabilities of growth rate at each temperature, means and 95% HPD intervals. D) Coefficients of genetic and environmental variation for each temperature, means and 95% HPD intervals. Note that points obscure small error bars. E) Principle component analysis of the **G**-matrix: proportions of variance explained by the different components. Error bars are 95% HPD intervals. F) Loadings of components 1 and 2 for each temperature. Error bars are 95% HPD intervals.

Most of the variation observed in growth rates was due to genetic variation present among the strains. Heritabilities for growth at different temperatures were high, around 0.89 for temperatures from 20 to 35 °C (Figure 3C). As temperature increased further heritability dropped to 0.63 at 40 °C (Figure 3C). However, this lower heritability was not due to decreased genetic variation but increased environmental variance at 37.5 and 40 °C (Table 1). Therefore there was substantial genetic variation for growth rate at 40 °C but environmental variation increased as well; looking at heritability alone would have been misleading in this case. Furthermore, as trait means differ across the different temperatures looking at genetic variances alone would have suggested that 35 °C has the most genetic variance (Table 1), but this would have been also misleading as coefficient of genetic variation reveals that growth at 40 °C has the most genetic variation followed by the other temperatures in decreasing order (Figure 3D). The same was true for coefficient of environmental variation (Figure 3D).

Eigen decomposition of the **G**-matrix can reveal what are the main axes along which correlated traits most readily evolve. We used principle component analysis to decompose the **G**-matrix. The first two principle components explained most of the variance with the first component explaining 79.5% (72.4%–86.0%) and the second component 19.3% (13.1%–26.6%) of the variance (Figure 3E). The rest of the components explained the remaining 1.2% of the variance, but the sizes of their corresponding eigenvalues were so small that this 1.2% is unlikely to have any biological meaning. Moreover, the interval estimates for the loadings of components 1 and 2 were consistent with no sign changes (Figure 3F), but this was not the case for rest of the components, indicating that loadings for the rest of the components are very uncertain. All the loadings of the first principle component were positive (Figure 3F), indicating that most variation in tolerance curves is mainly for elevation. The second component suggested that growth rate at 40 °C and to lesser extent at 37.5 °C are more independent from rest of the temperatures, even though some variation is shared with 40 °C and the rest of the temperatures, as genetic correlation with 40 °C and the other temperatures were positive (Table 1).

When looking trait specific evolvabilities we also observed that growth rate at 40 °C had the highest conditional evolvability and the highest autonomy (Table 3). This indicates that out of all of the growth rates, growth rate at 40 °C can evolve by itself most easily. The rest of the traits had very low autonomies reflecting their high genetic correlations (Tables 1 and 3).

**Table 3:**
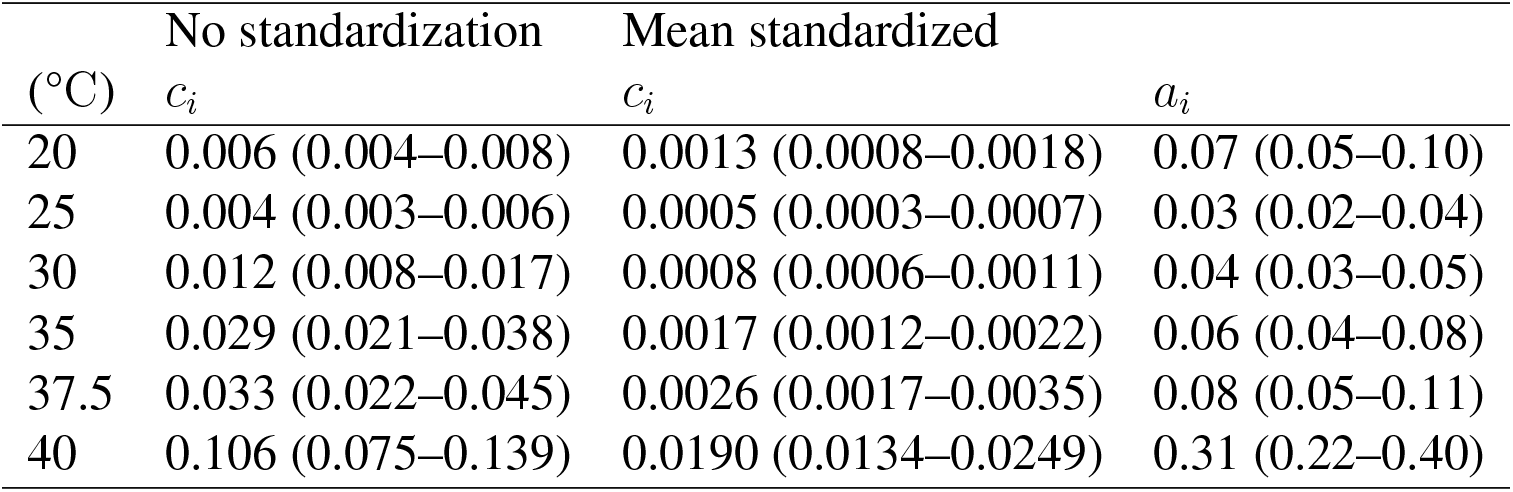
Conditional evolvabilities (*c_i_*) and autonomies (*a_i_*) for growth rates at different temperatures, values are posterior medians and 95% HPD interval is shown in parenthesis. For conditional evolvability values for both without standardization and with mean standardized **G**-matrices are shown. Values for trait specific autonomy are the same with and without standardization.

| (°C) | No standardization | Mean standardized |  |
| --- | --- | --- | --- |
| | $c_i$ | $c_i$ | $a_i$ |
| 20 | 0.006 (0.004–0.008) | 0.0013 (0.0008–0.0018) | 0.07 (0.05–0.10) |
| 25 | 0.004 (0.003–0.006) | 0.0005 (0.0003–0.0007) | 0.03 (0.02–0.04) |
| 30 | 0.012 (0.008–0.017) | 0.0008 (0.0006–0.0011) | 0.04 (0.03–0.05) |
| 35 | 0.029 (0.021–0.038) | 0.0017 (0.0012–0.0022) | 0.06 (0.04–0.08) |
| 37.5 | 0.033 (0.022–0.045) | 0.0026 (0.0017–0.0035) | 0.08 (0.05–0.11) |
| 40 | 0.106 (0.075–0.139) | 0.0190 (0.0134–0.0249) | 0.31 (0.22–0.40) |

### Evolution of performance curves

In order to examine how a performance curve of a population that has the same **G**-matrix as estimated here could evolve, we performed simulations with a quantitative genetic model of performance curve evolution. First we asked how performance curves responded to selection if selection were to operate in the same direction as the two first observed loadings of the **G**-matrix eigen decomposition (Figure 3F). We normalized the summed absolute values of selection gradients across all temperatures to be 0.6 mm/h and their relative weights to be proportional to the loadings of each principle component. Theoretical prediction is that when ***β*** is in the same direction as the first component, evolvability should be the greatest (Schluter, 1996). Indeed, this is what we observed, as consequently response to selection was also greatest in this direction (Figure 4). Moreover, evolvability and conditional evolvability greatly surpassed the average evolvability across the entire phenotypic space (Figure 4D). When selection gradient pointed to the direction of the second component, unconditional evolvability was no longer larger than expected, while conditional evolvability still remained larger than average (Figure 4D).

**Figure 4:**
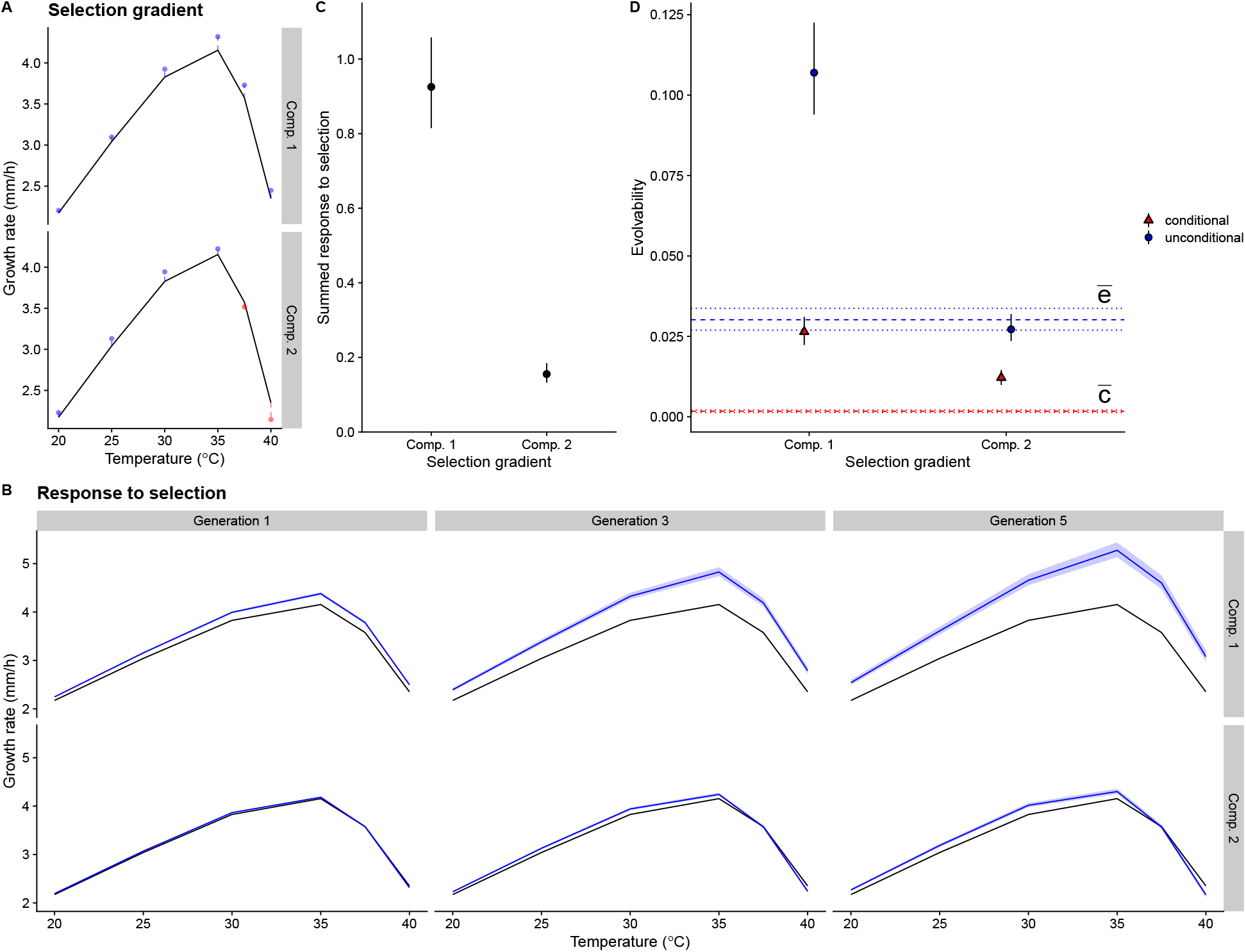
Simulated responses to selection using selection gradient, ***β***. A) Selection gradients correspond to the loadings of the first two components of the **G**-matrix eigen decomposition. Black line is the mean empirical performance curve and dots represent values of selection gradient for each temperature. Blue dots represent selection for increased growth and red dots for decreased growth. B) Simulated responses to selection, black line is the empirical mean and blue lines are the simulated performance curves after selection. Shaded regions contain 95% of the simulations. Note that variability due to uncertainty in the **G**-matrix is not visible for many of the simulations. Columns show selection responses after 1, 3, or 5 generations of selection and rows show results for different selection regimes. C) Summed absolute values for response to selection in a single generation for the two gradients. D) Medians and 95% intervals for mean standardized evolvabilities for the two gradients, blue horizontal lines (*ē*) show the average unconditional evolvability across random selection gradients and red horizontal lines (*c̄*) show the average conditional evolvability, dotted lines show the 95% HPD interval.

Next we examined responses to different selection differentials with the idea that we want to know whether particular phenotypic change in the performance curve was possible, whatever the selection gradient implied by the selection differentials. For instance, when we simulated selection for increased growth at a single temperature this leads to positive correlated responses in other temperatures if the other traits are neutral, as in the case when selection gradient is zero for a given trait. However, when there was selection for increased growth at a single temperature and to maintain the original phenotype at the other temperatures there were also correlated responses but these were less uniform (Figure S3). Accordingly, selection at a single temperature often lead to correlated responses in nearby temperatures (Figure S4). Selection at multiple temperatures lead to stronger responses to selection and correlated responses (Figures S5 and S6). For instance, selection at 25 and 30 °C increased growth rate also at 20 °C (Figure S5). When selection happened at multiple temperatures, response could be bigger in certain temperature than if selection happened for that temperature alone. For example, if there was selection for higher growth at 20, 25, and 30 °C, response to selection was greater than if there was selection for higher growth only at 20 °C (Figure S4 and S6). With selection differential of 0.2 only at 20 °C, response to selection after five generations was 2.80 (2.75–2.84, 95% HPD). Whereas if selection differential was 0.2 at 20, 25, and 30 °C, response to selection after 5 generations of selection was 3.01 (2.98–3.04, 95% HPD). Thus, it was not possible to change a certain temperature completely independently of the others, but often extreme temperatures could be changed without affecting the growth at the other extreme. We then asked is it possible to create similar evolutionary responses in performance curves as shown in Figure 1. We were able to find a set of selection differentials that were able to generate changes in elevation, horizontal shift, or shape (Figure 5). This shows that despite strong genetic correlations it is possible for the performance curves to evolve in almost any manner if selection favors such a performance curve. However, selection regimes involving horizontal shifts require selection for increased growth rate in some temperatures and decreased growth rate in others (Figure 5). Evolvabilities and conditional evolvabilities were highest for elevation changes. For optimum shifts and shape changes conditional evolvabilites were lower than the average conditional evolvability over all phenotypic space (Figure 5C). This indicates that elevation changes are less constrained than changes in optimum temperature or performance curve shape.

**Figure 5:**
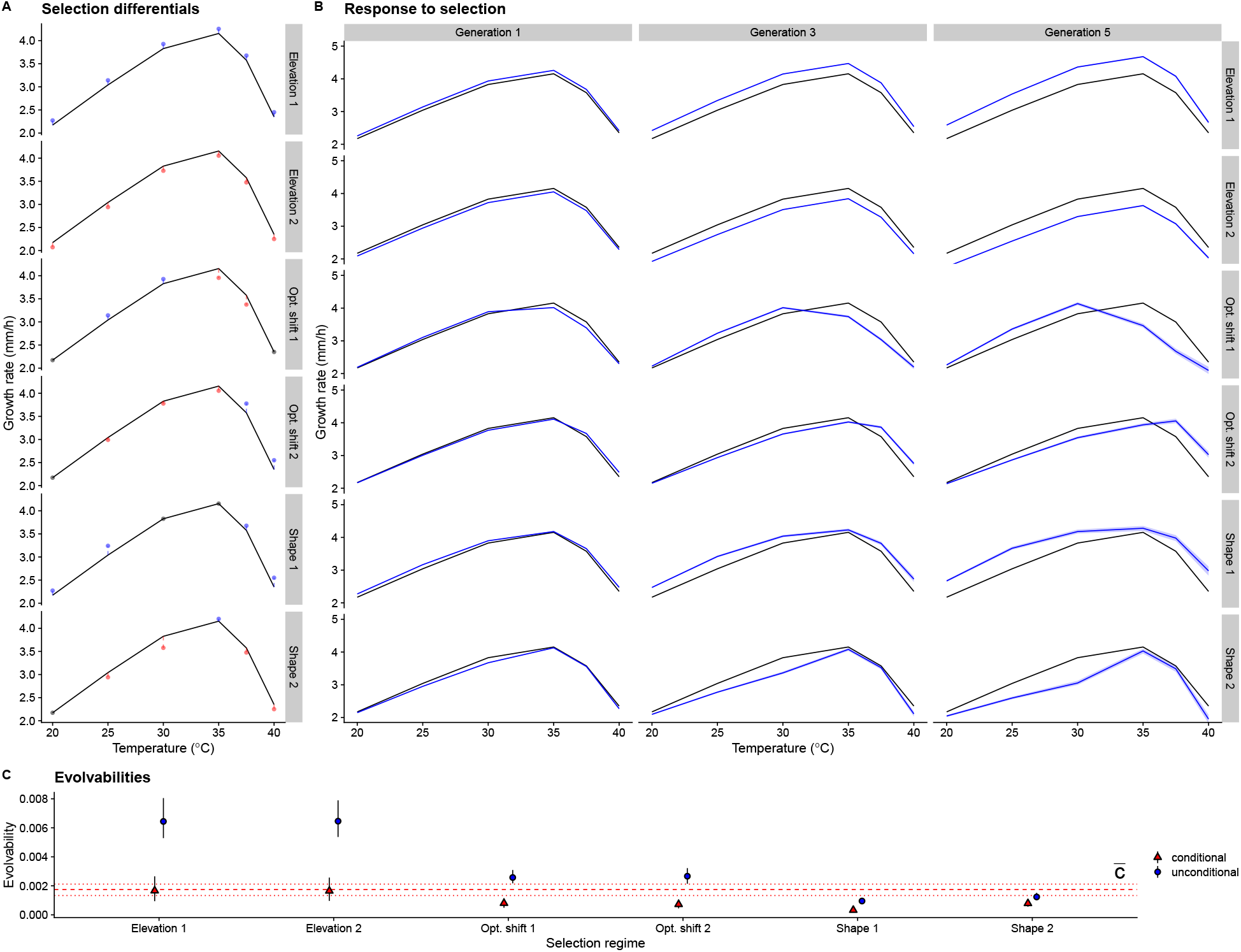
Simulated responses to selection using selection differentials, **S**. A) Selection differentials for each selection regime. Black line is the mean empirical performance curve and dots represent values of selection differentials for each temperature. Blue dots represent selection for increased growth, red for decreased growth and black dots indicate stabilizing selection at this temperature. Selection regime 1 selects for increased elevation, regime 2 selects for decreased elevation, regime 3 selects for lower optimum temperature, regime 4 selects for higher optimum, regime 5 selects for broader shape, and regime 6 selects for narrower shape. B) Simulated responses to selection. C) Evolvability and conditional evolvability for each of the selection gradients implied by the selection differentials. Red horizontal lines show the average conditional evolvability (*c̄*) across the entire phenotypic space. Average evolvability is higher than the y-axis scale and is not shown.

## Discussion

We have shown that there is substantial genetic variation in thermal performance curves of *Neurospora crassa*. Most of this variation is in performance curve elevation and there is very little evidence of strong trade-offs. Genetic variation in growth is strongly correlated among nearby temperatures but there is a threshold before or at 40 °C after which this correlation drops, indicating that physiological processes at 40 °C are different than those at lower temperatures. Such thresholds are common in many organisms, including *Drosophila* where different expression profiles were observed in cold, moderate, and hot temperatures (Colinet et al., 2013).

In many ways, variation in performance curves of *N. crassa* are quite typical for many ectotherms that have been studied (Sinclair et al., 2016). Most genetic variation in *N. crassa* is variation in performance curve elevation, which contrasts with previous studies in other species that have found most variation to be for reaction norm shapes (Izem and Kingsolver, 2005; Logan et al., 2020). Yet variation in performance curve elevation is commonly found, a review of thermal performance curves in insects found that elevation shifts were the most common type of change along environmental gradients (Tüzün and Stoks, 2018), see also Scheiner (1993). We also observed quite substantial genetic variation in thermal performance, and while comparisons between animals and fungi should be treated with caution, other studies have observed much lower heritabilities (e.g. Logan et al., 2018; Castañeda et al., 2019; Martins et al., 2019).

Genetic variation in performance curve elevation could reflect differences in genetic condition of individuals, rather than temperature specific adaptation. This could be due to different strains harboring different amounts of deleterious mutations. However, this seems an unlikely explanation as *N. crassa* is haploid, so deleterious mutations are immediately exposed to selection and would be removed, as in nature there is plenty of sexual reproduction as indicated by rapid decay of linkage disequilibrium in the population genetic data (Ellison et al., 2011; Palma-Guerrero et al., 2013). Another possibility is that genetic differences between the strains in how well they are able to grow in lab conditions are thermodynamically amplified, as increasing temperature also increases metabolic rate (Schulte, 2015). However, our estimates of activation energy were much lower than the thermodynamic expectation, and contrast with previous studies that have found much stronger relationship between growth rate and optimum temperatures (Savage et al., 2004; Knies et al., 2009; Sørensen et al., 2018). While we cannot exclude that some of the differences were due to the thermodynamic effect, this cannot be the whole explanation as there were clear genotype by environment interactions indicated by genetic correlations across environments that were less than one. There have to be alleles segregating in the population that have different effects in different temperatures. Particularly, genetic variation after the optimum of the thermal performance curve has been reached cannot be accounted by thermodynamic effects (Schulte, 2015).

There was no indication of strong trade-offs between temperatures, and certainly not the kind of trade-offs that have been assumed in many models of tolerance curve or reaction norm evolution in general (Angilletta et al., 2003). The absence of any trade-offs suggests that theoretical models of reaction norm evolution that assume trade-offs should be treated with caution. It further poses a question: if growth rate is closely linked to fitness, and if there are no trade-offs, why there is genetic variation in growth? It seems reasonable that mycelial growth rate should be a fitness component in filamentous fungi. In a previous study no trade-off was detected between mycelial growth rate and spore production (Anderson et al., 2018). However, there is some evidence that strains that have higher growth rates have also higher competitive fitness (Kronholm et al., 2020). It may be that there is a trade-off between growth rate and some other trait which we have not measured, for example Ketola et al. (2013) found a trade-off between bacterial virulence and growth in high temperatures. Alternatively, the evolution of performance curves may be limited by the environments, and thus the selection pressures, the strains encounter rather than genetic tradeoffs (Whitlock, 1996; Kassen, 2002). If there is no selection at a particular temperature, then variation at those temperatures may be neutral. The evidence for trade-offs and cost of plasticity for temperatures has been mixed; some studies have observed trade-offs (Knies et al., 2006; Romero-Olivares et al., 2015; Le Vinh Thuy et al., 2016), while others have observed that adapting to one temperature did not limit plasticity (Fragata et al., 2016; Manenti et al., 2015), most genetic variation has been observed for overall performance (Klepsatel et al., 2013; Latimer et al., 2015), or that adaptation was largely temperature specific with no apparent trade-offs (Bennett et al., 1992). Genetic correlations between growth rates at nearby temperatures were strong, which is to be expected, as a difference of a few °C is likely to be a very similar physical environment for an organism. However, growth rate at 40 °C had a lower genetic correlation to growth rates at other temperatures. This suggests that at 40 °C there was some physiological process activated, which has genetic variation, but that was not active or was at much lower level of activity in lower temperatures. The most obvious candidate for such a process is the heat shock response (Piper, 1993; Feder and Hofmann, 1999; Sørensen et al., 2003). Previously the heat shock response of *N. crassa* has been studied at 42 °C or higher (Mohsenzadeh et al., 1998; Plesofsky-Vig and Brambl, 1985; Guy et al., 1986) but it probably occurs already at lower temperatures, as we observed significant slow down of growth at 40 °C. The canonical heat shock proteins are important for the physiological heat shock response, but there can be additional mechanisms involved: there is evidence that the sugar trehalose plays some role in *N. crassa* heat shock response (Bonini et al., 1995). Furthermore, changes in cell membrane composition are involved in temperature acclimation and the proportion of highly unsaturated fats increases in low temperatures (Martin et al., 1981). These responses have been observed in yeasts as well (Glatz et al., 2015). It is likely that there is genetic variation in the heat shock response induction threshold or in the magnitude of heat shock response, and this physiological variation can explain why genetic correlation across temperatures is lower when 40 °C is involved. Further investigation into variation of heat shock responses at the physiological level seems warranted.

## Conclusions

At the species level, populations of *N. crassa* contain plenty of genetic variation for growth at different temperatures, and may be able to respond to increasing temperatures and thermal fluctuations via genetic adaptation mainly by increasing overall performance. An experimental evolution study with a related species, *N. discreta*, also found adaptation to higher temperature (Romero-Olivares et al., 2015). Previous studies have suggested that warming may pose the greatest risk to tropical animal species, as they live already close to their thermal maxima (Deutsch et al., 2008), but *N. crassa* is different in this respect. Whether this is true for all fungi or if *N. crassa* is a special case remains to be investigated.

We did not observe any inherent genetic trade-off between hotter and colder temperatures, which is in contrast to common theoretical assumptions. Thermal performance curves of *N. crassa* can in theory evolve to have nearly any shape provided that appropriate selection gradient exists. Whether such selection gradients occur in nature is another matter. However, it seems more plausible that if there would be selection for increased growth at higher temperatures, evolutionary response will happen by either increasing the overall elevation of the performance curve, which was the line of least genetic resistance.

Revealing the genetic basis of performance curve variation is a topic for future studies, and would allow investigating whether trade-offs exists at the level of specific alleles. We are pursuing this question in future work.

## Acknowledgements

This study was funded by grants from Emil Aaltonen foundation and Ella & Georg Ehrnrooth foundation to IK and Academy of Finland Research Fellowships to IK (no. 321584) and TK (no. 278751). We’d like to thank Matthieu Bruneaux for comments on the manuscript.

## Supplementary Information

### Supplementary Figures

**Figure S1:**
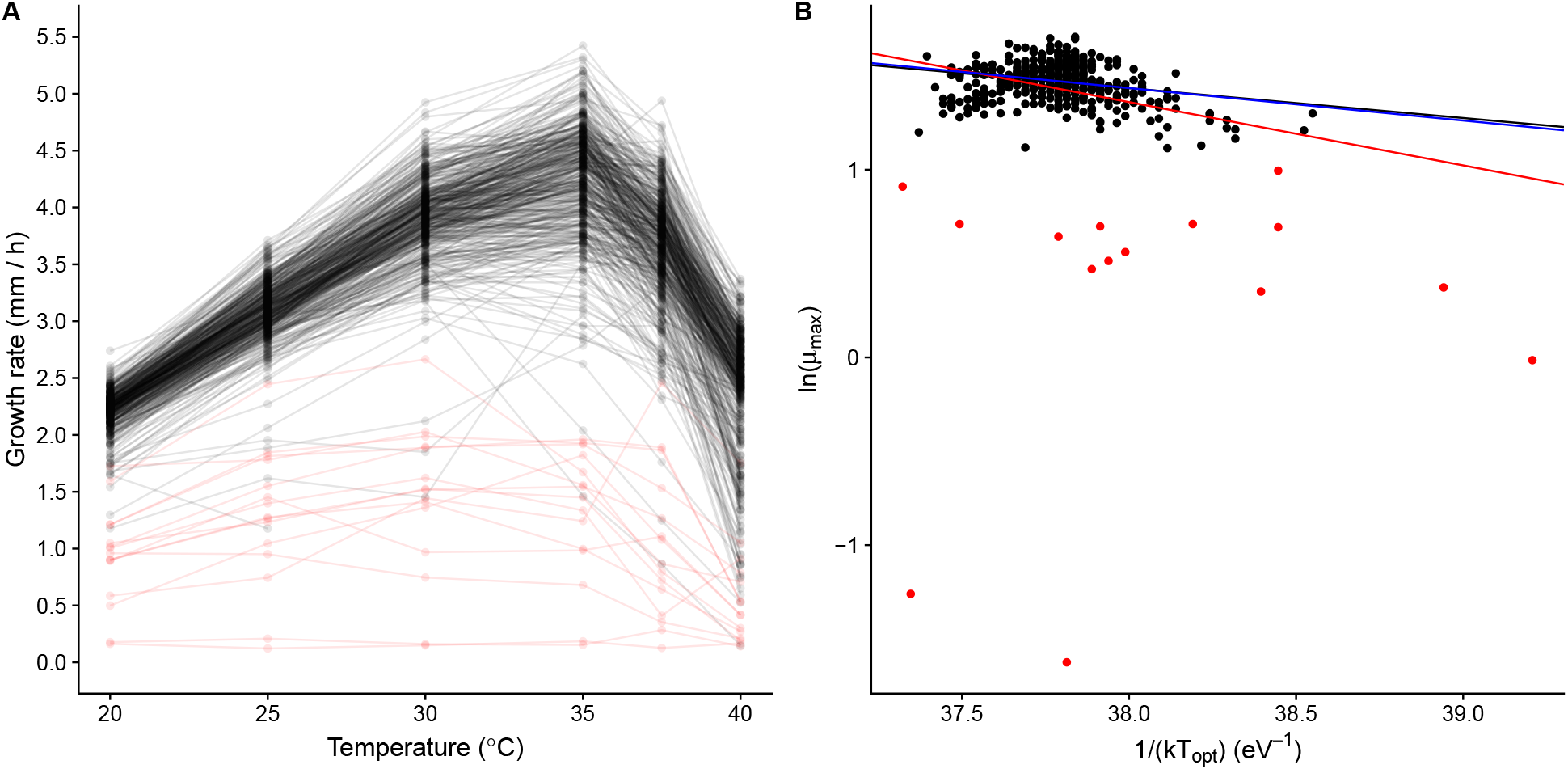
A) Phenotypic means for each genotype, those genotypes that were removed from the thermodynamic analysis as outliers are coloured red. B) Logarithm of maximum growth rate, *μ_max_*, plotted against inverse of *kT_opt_*. Datapoints that were removed as outliers are coloured red. Black regression line is ordinary regression fitted to the data without outliers (red points removed), slope (±SE) is −0.16(±0.03). Red line is ordinary regression fitted to all of the data, slope is −0.34(±0.06). Blue line is robust regression with bisquare weights fitted to all of the data, slope is −0.17(±0.03).

**Figure S2:**
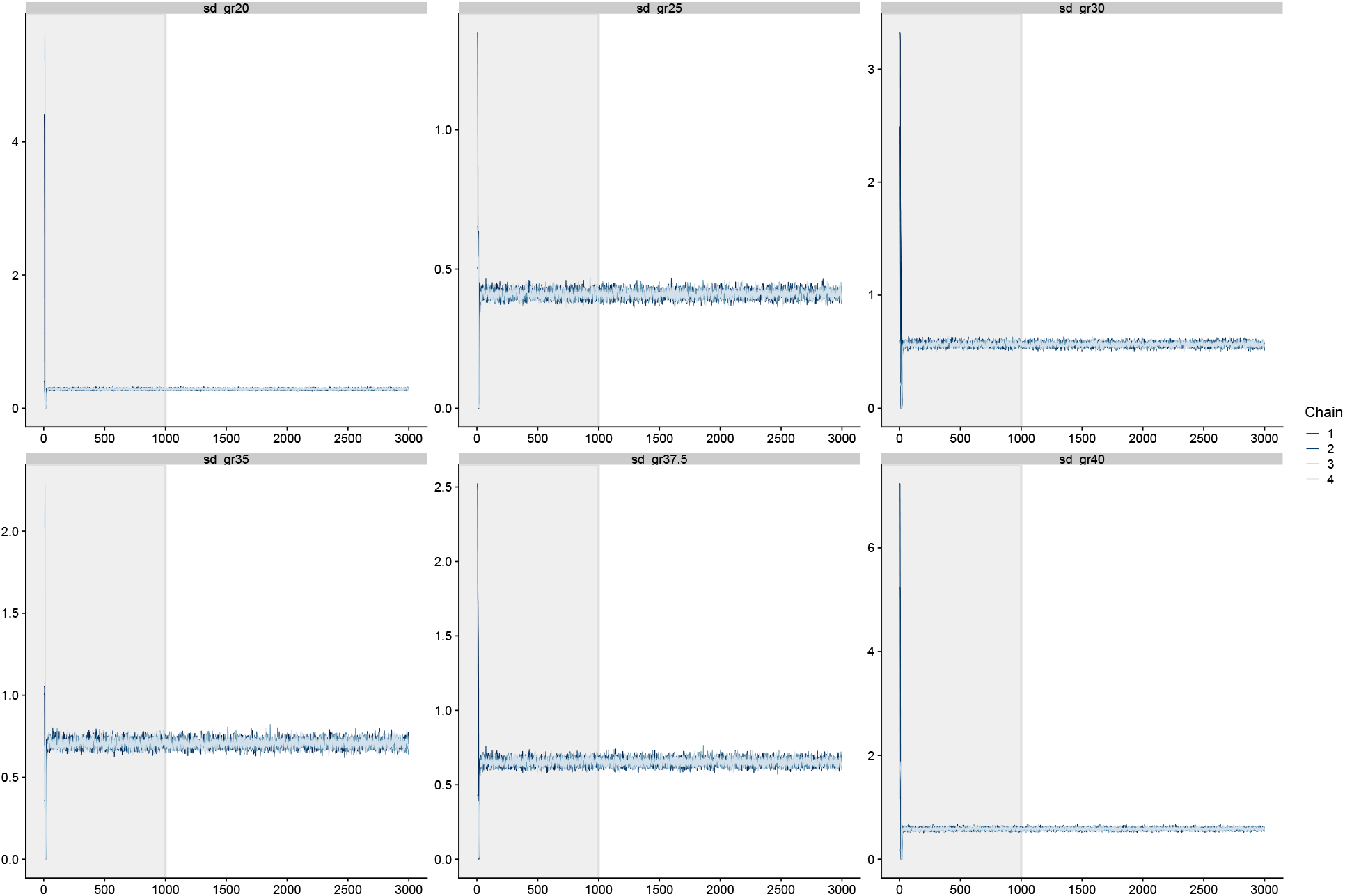
Example MCMC traceplots for genetic standard deviations of growth rates in different temperatures in the multivariate model. The grey shaded area denotes the warmup iterations which were discarded from the final parameter estimates. In this example four independent chains were initialized at random values; the chains rapidly converge to the same distribution during warmup. No divergent transitions were observed in this run.

**Figure S3:**
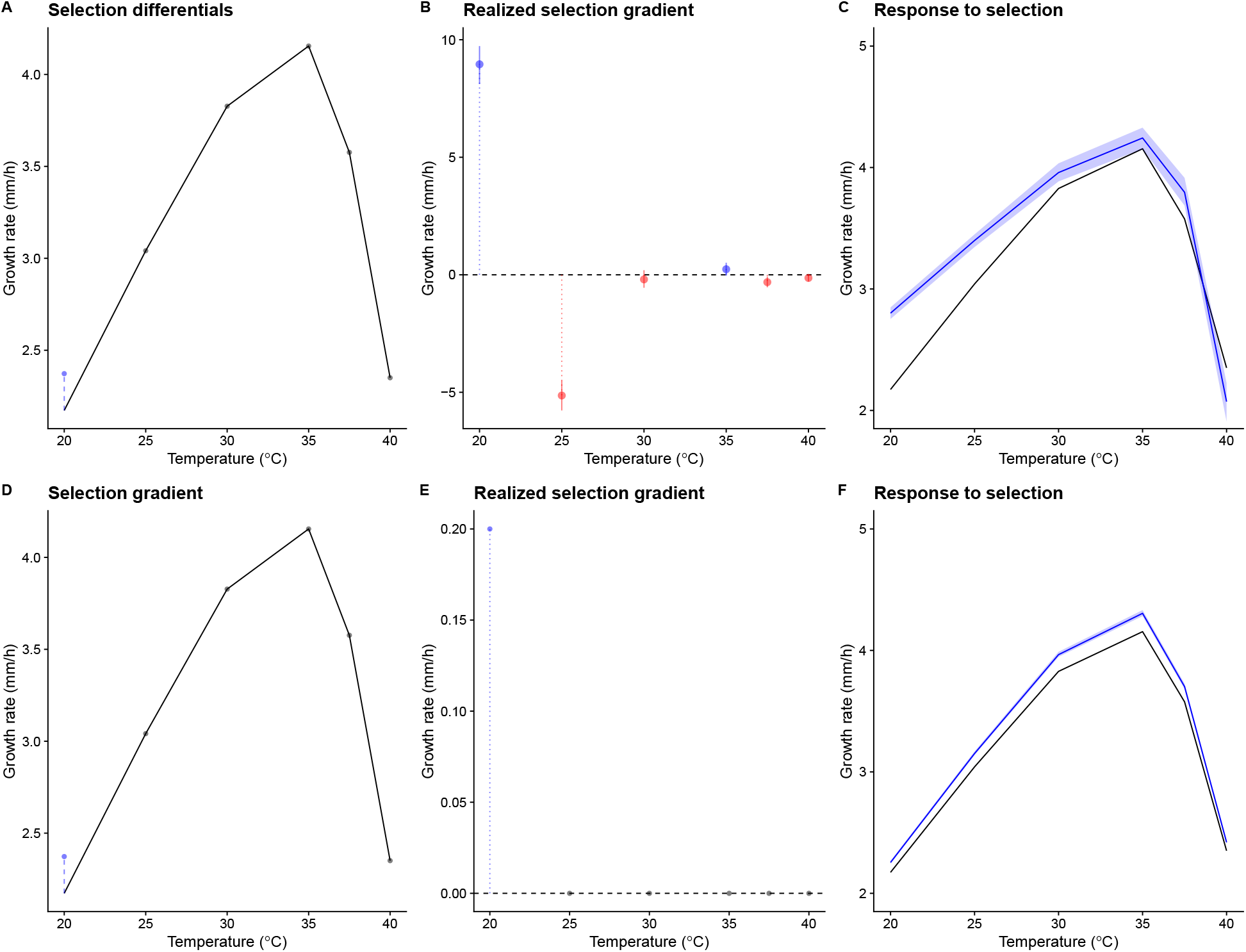
Illustration on how selection differential and selection gradient generate different responses to selection. Top row: selection based on selection differentials. A) There is selection for increased growth at at 20 °C and to maintain the original phenotype for the other traits. Black line is the empirical performance curve and blue dots represent selection differentials for each temperature, **S** = 0.2, 0, 0, 0, 0, 0. B) The realized selection gradient (***β*** = **P**^−1^**S**) implied by this selection differential. C) Blue line shows the mean phenotype after 5 generations of selection, shaded area contains 95% of the simulations. Bottom row: selection based on selection gradient. D) There is selection for increased growth at 20 °C as in A) but selection gradient is ***β*** = 0.2, 0, 0, 0, 0, 0. E) Now realized selection gradient is the same as in (D), there is selection for increased growth in one temperature but phenotypes of other temperatures are selectively neutral. E) Response to selection after 5 generations of selection as in (C) for this selection gradient. Selection using similar selection differentials and selection gradient leads to different phenotypic responses.

**Figure S4:**
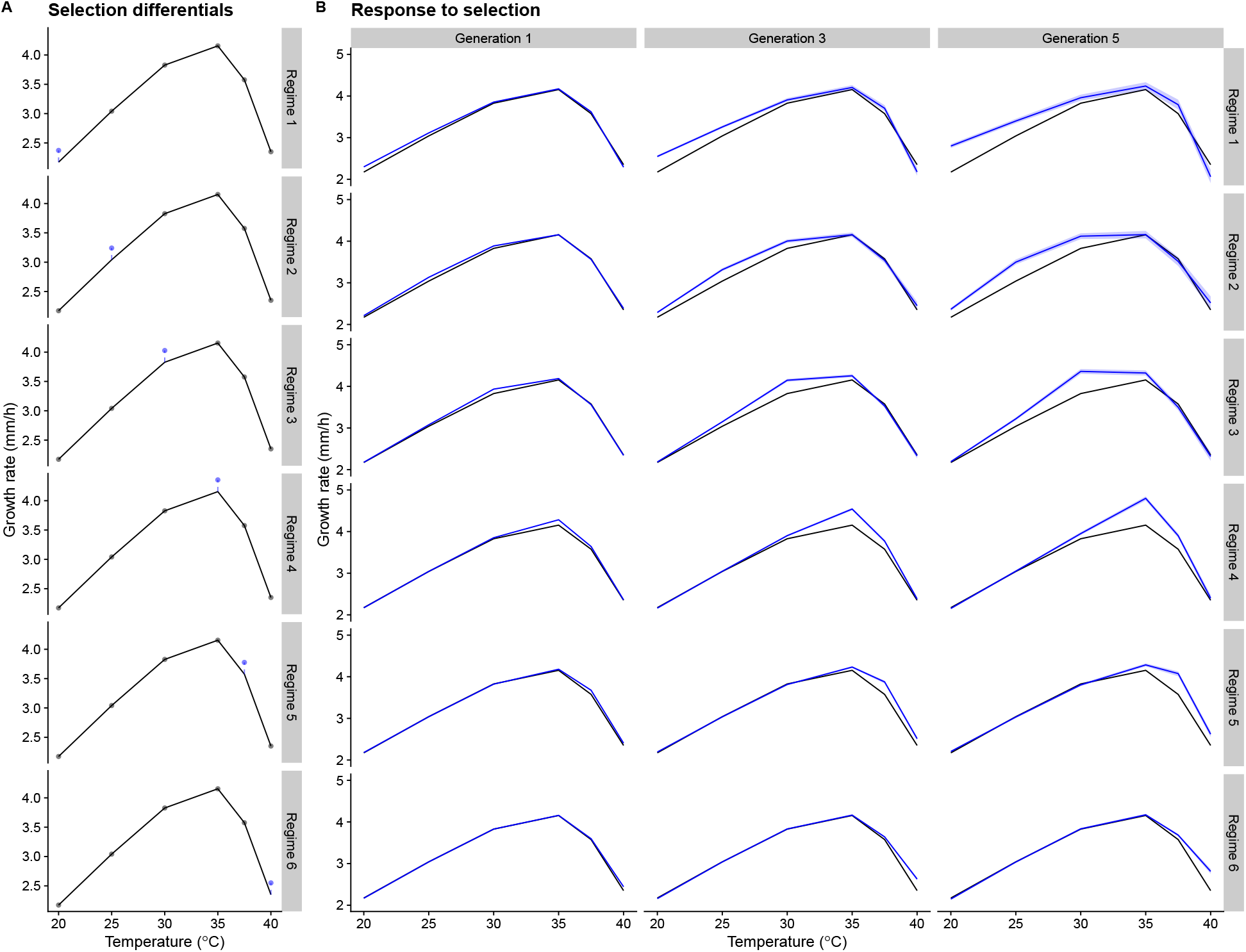
Selection for increased growth rate in a single temperature. Selection differential is 0.2 mm/h at each generation. A) Selection differentials for each selection regime. B) Simulated responses to selection.

**Figure S5:**
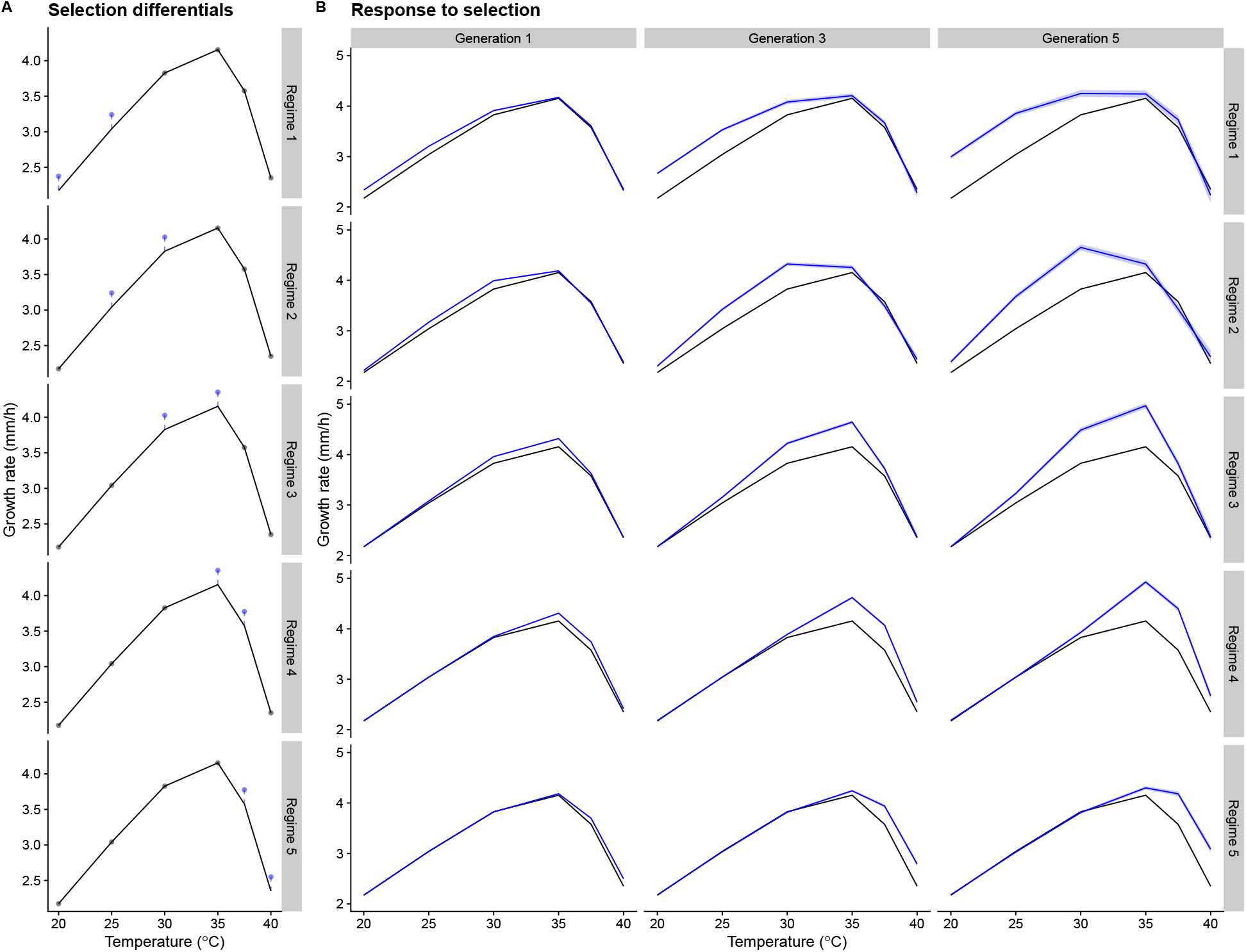
Selection for increased growth rate in two temperatures. Selection differential is 0.2 mm/h at each generation for each temperature, so 0.4 in total for each selection regime. A) Selection differentials for each selection regime. B) Simulated responses to selection.

**Figure S6:**
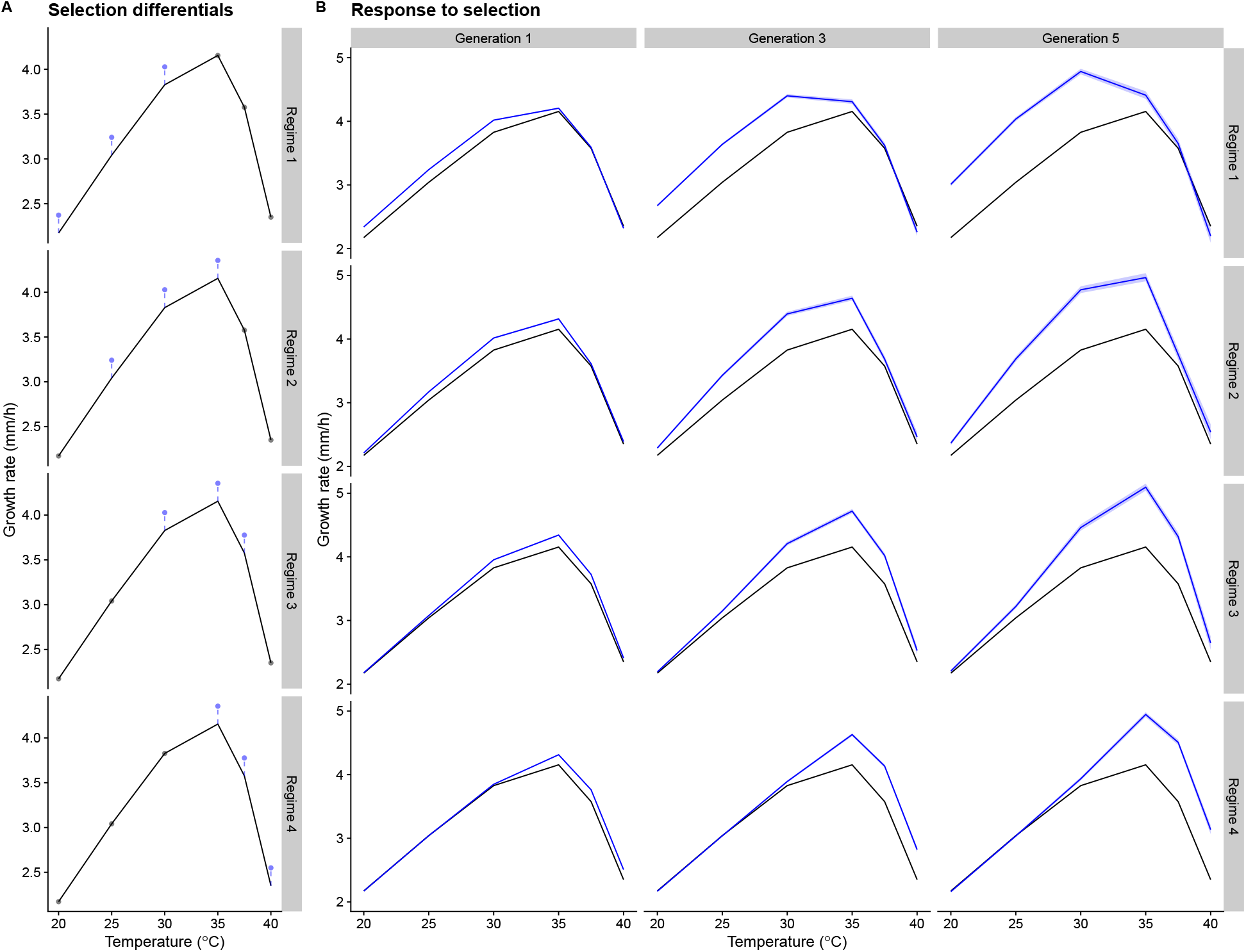
Selection for increased growth rate in three temperatures. Selection differential is 0.2 mm/h at each generation for each temperature, so 0.6 in total for each selection regime. A) Selection differentials for each selection regime. B) Simulated responses to selection.

### Supplementary Tables

**Table S1:**
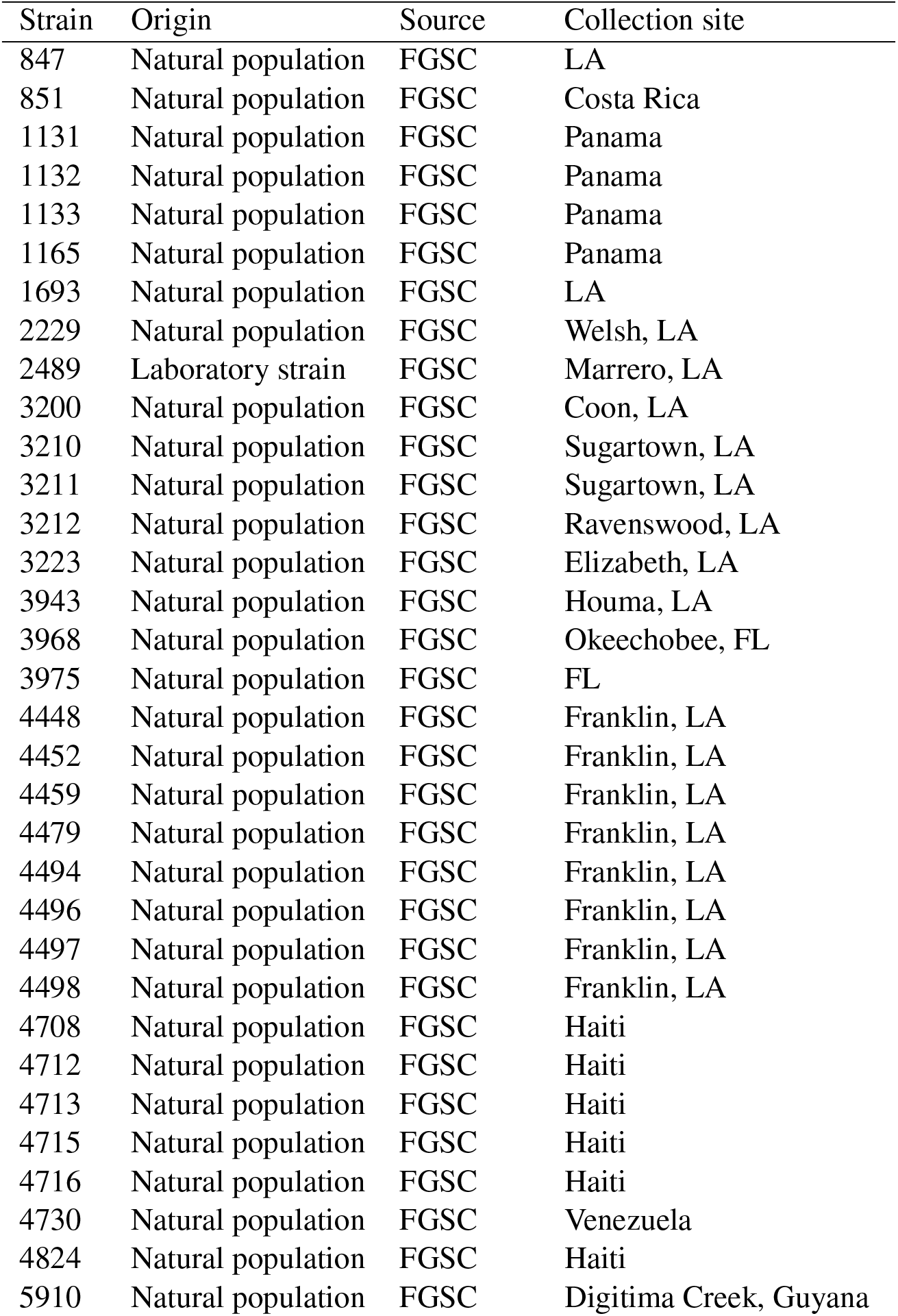
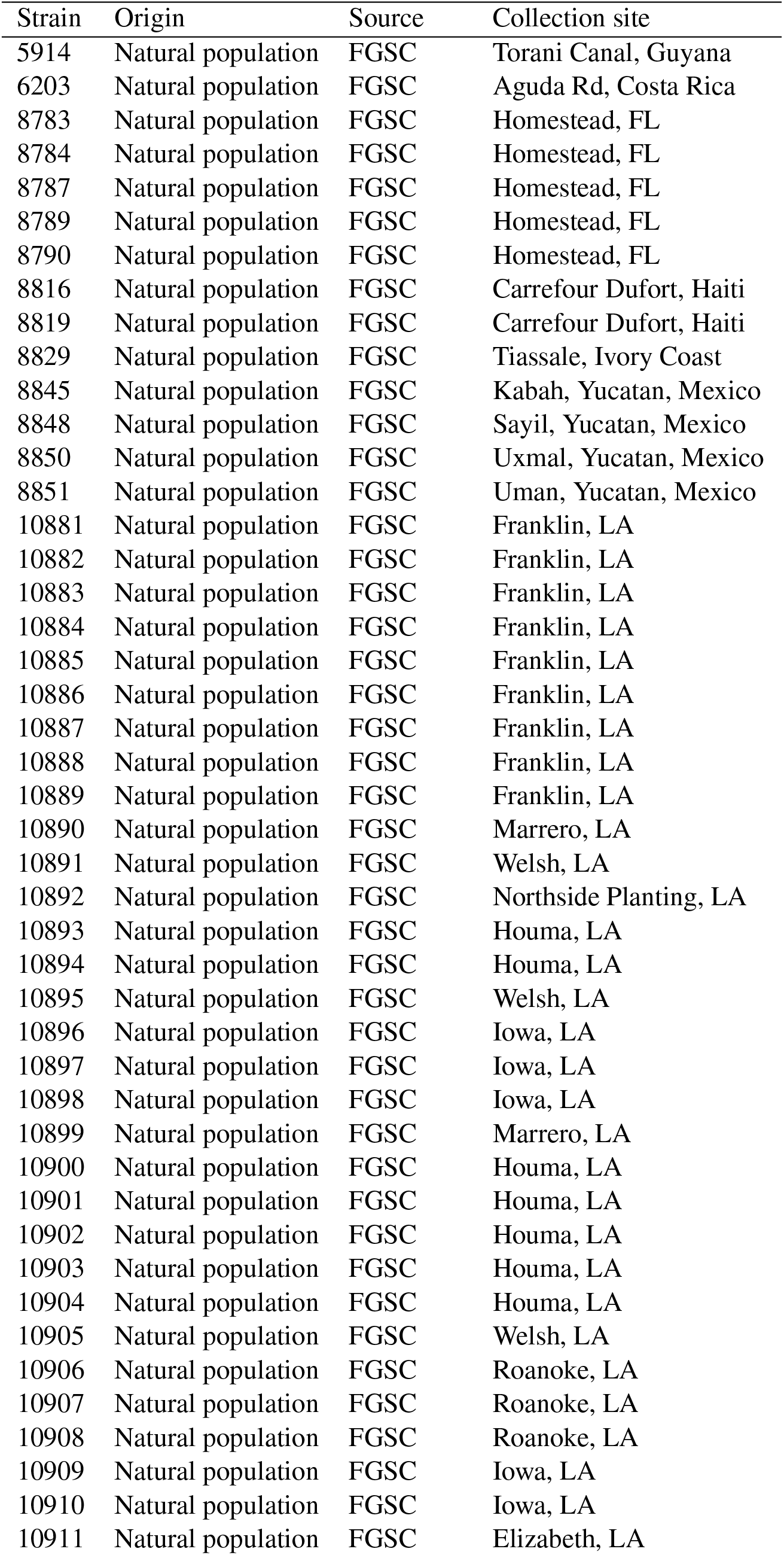
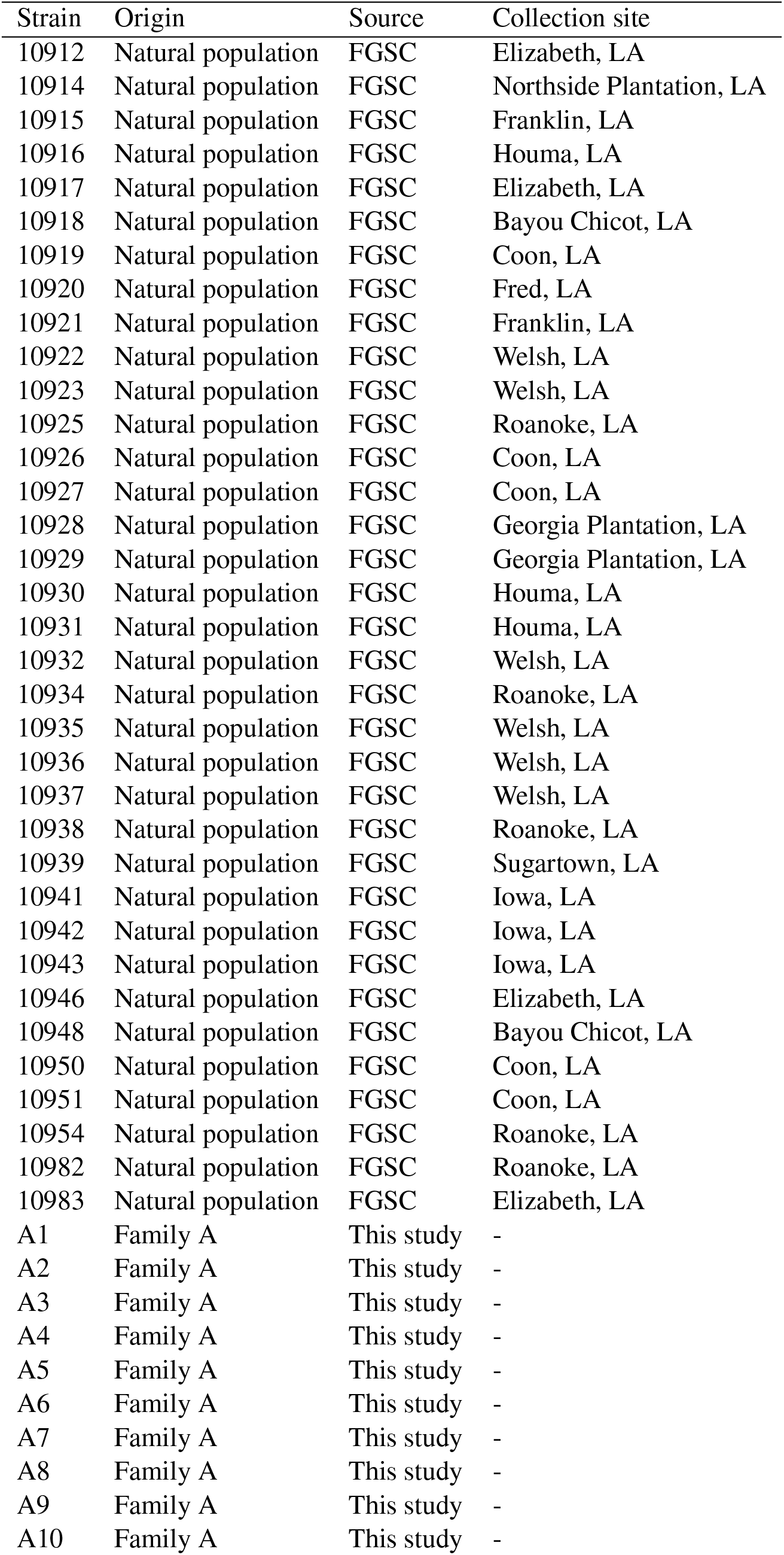
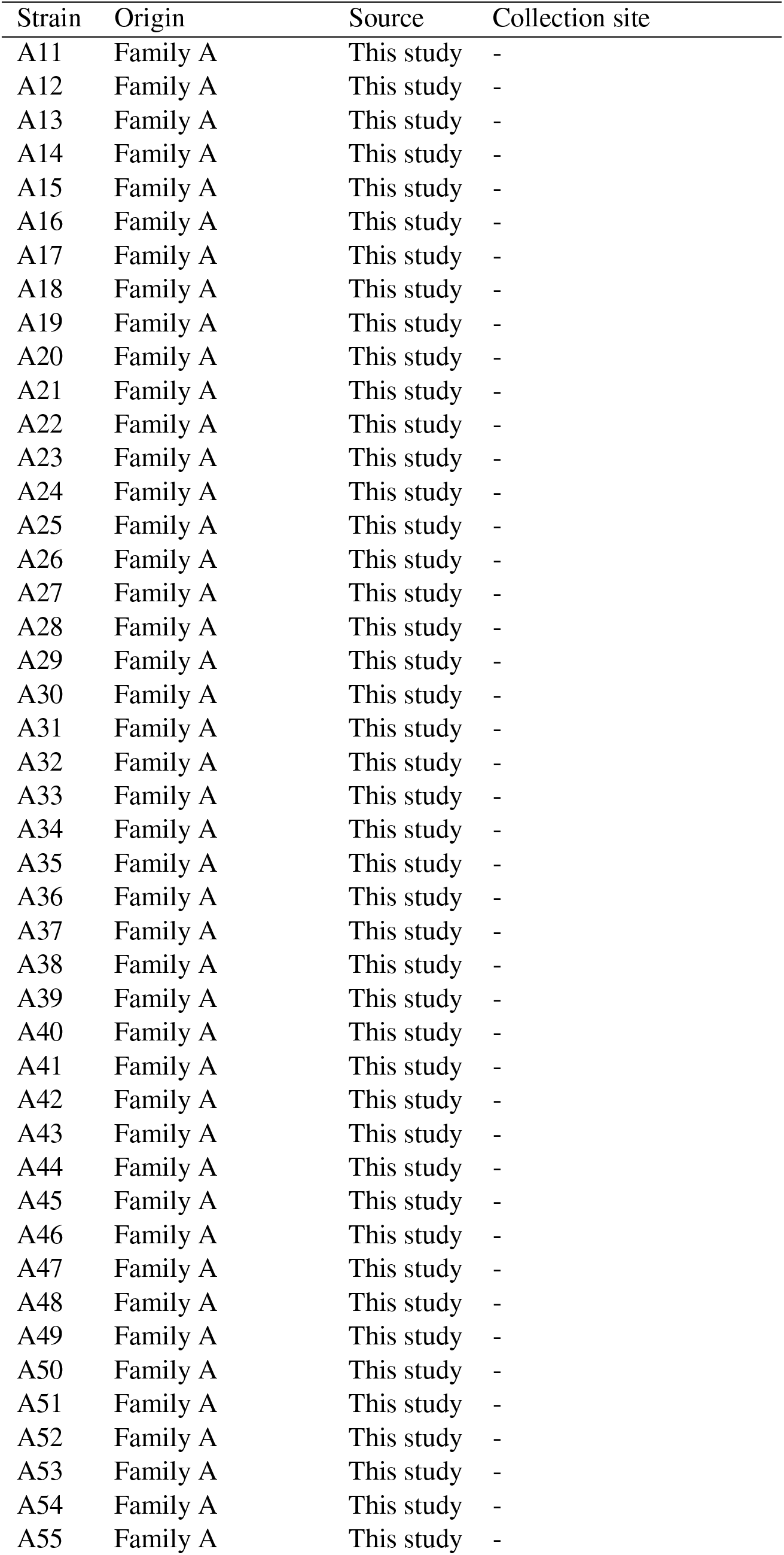
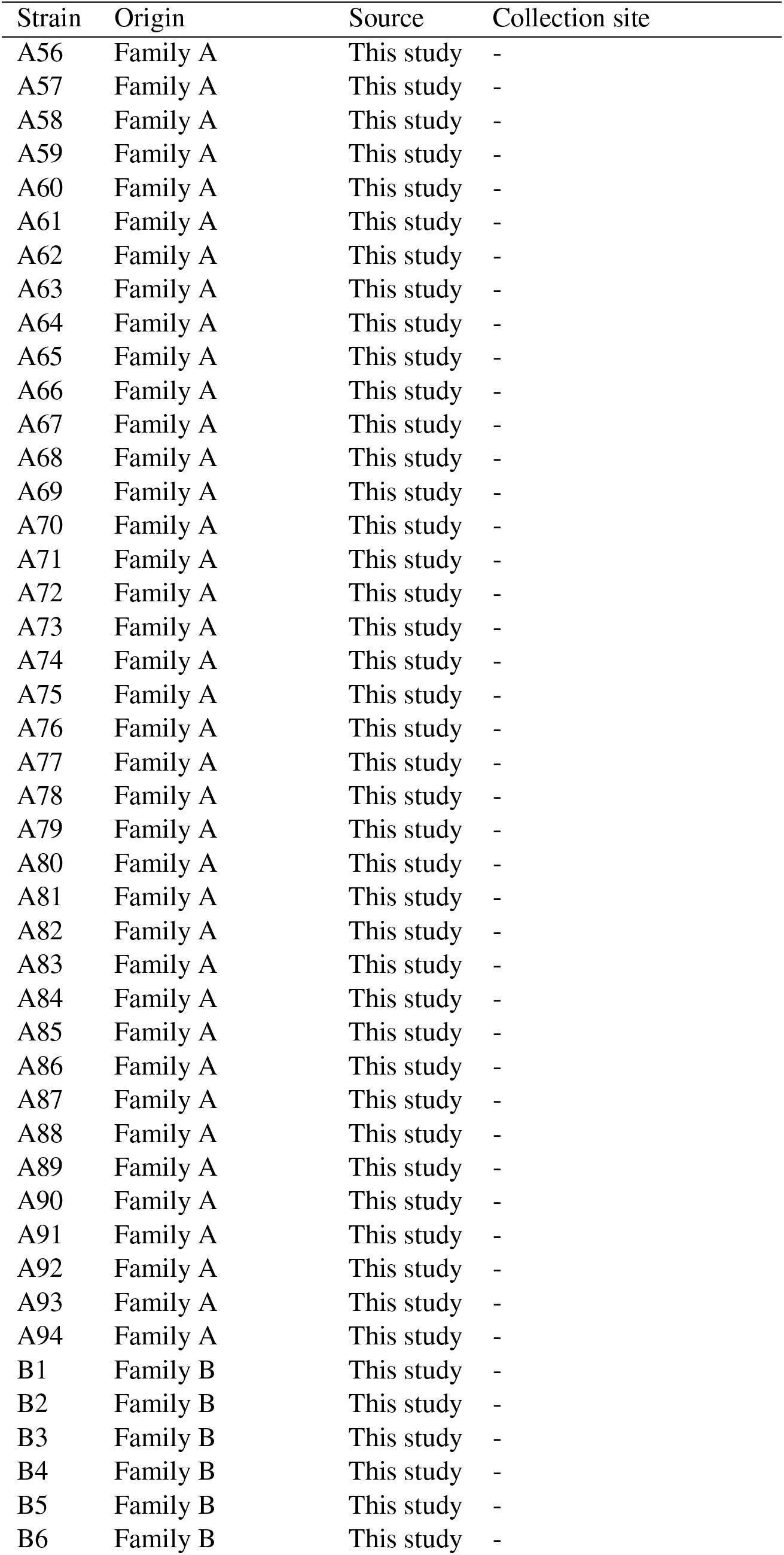
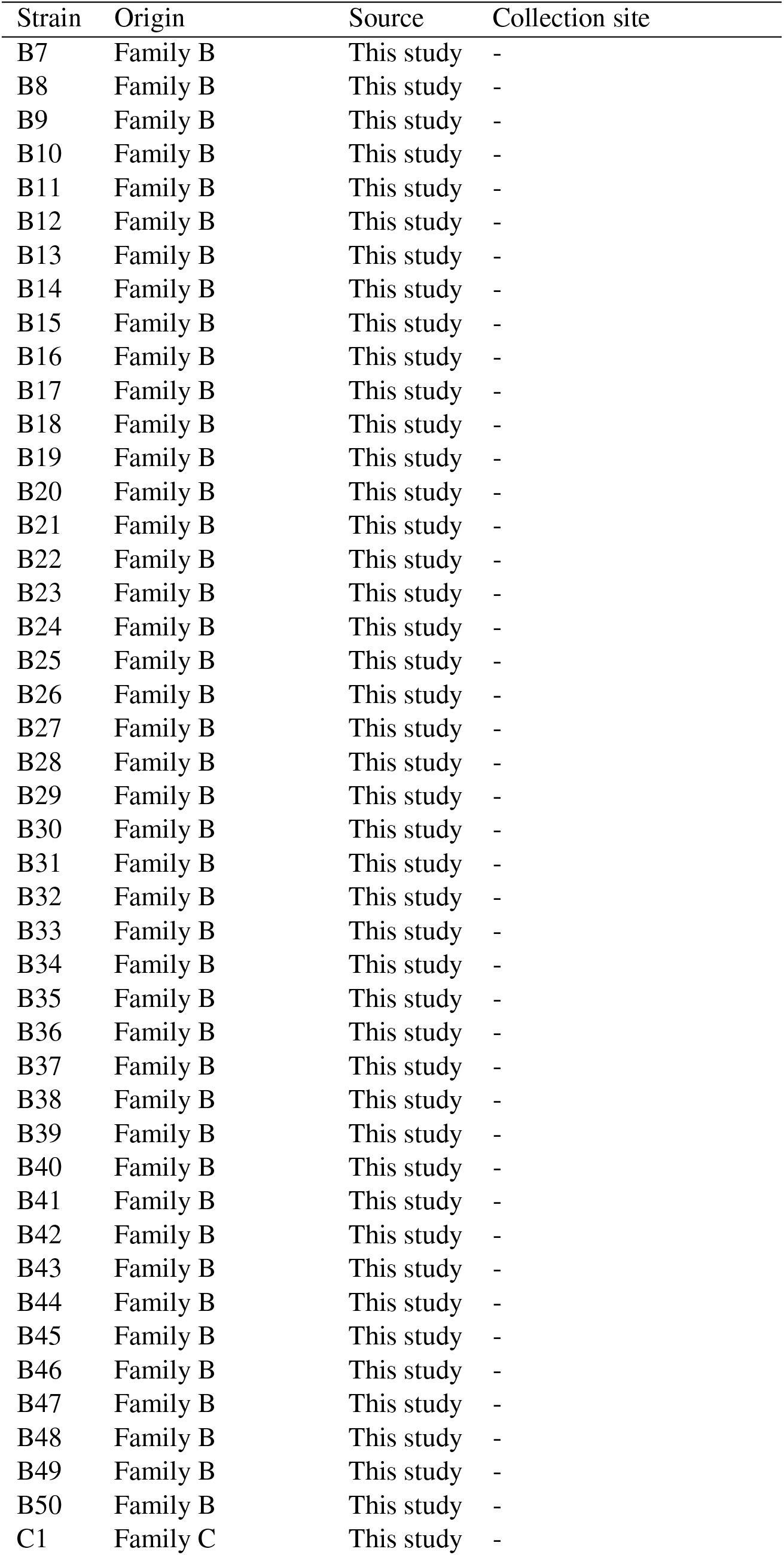
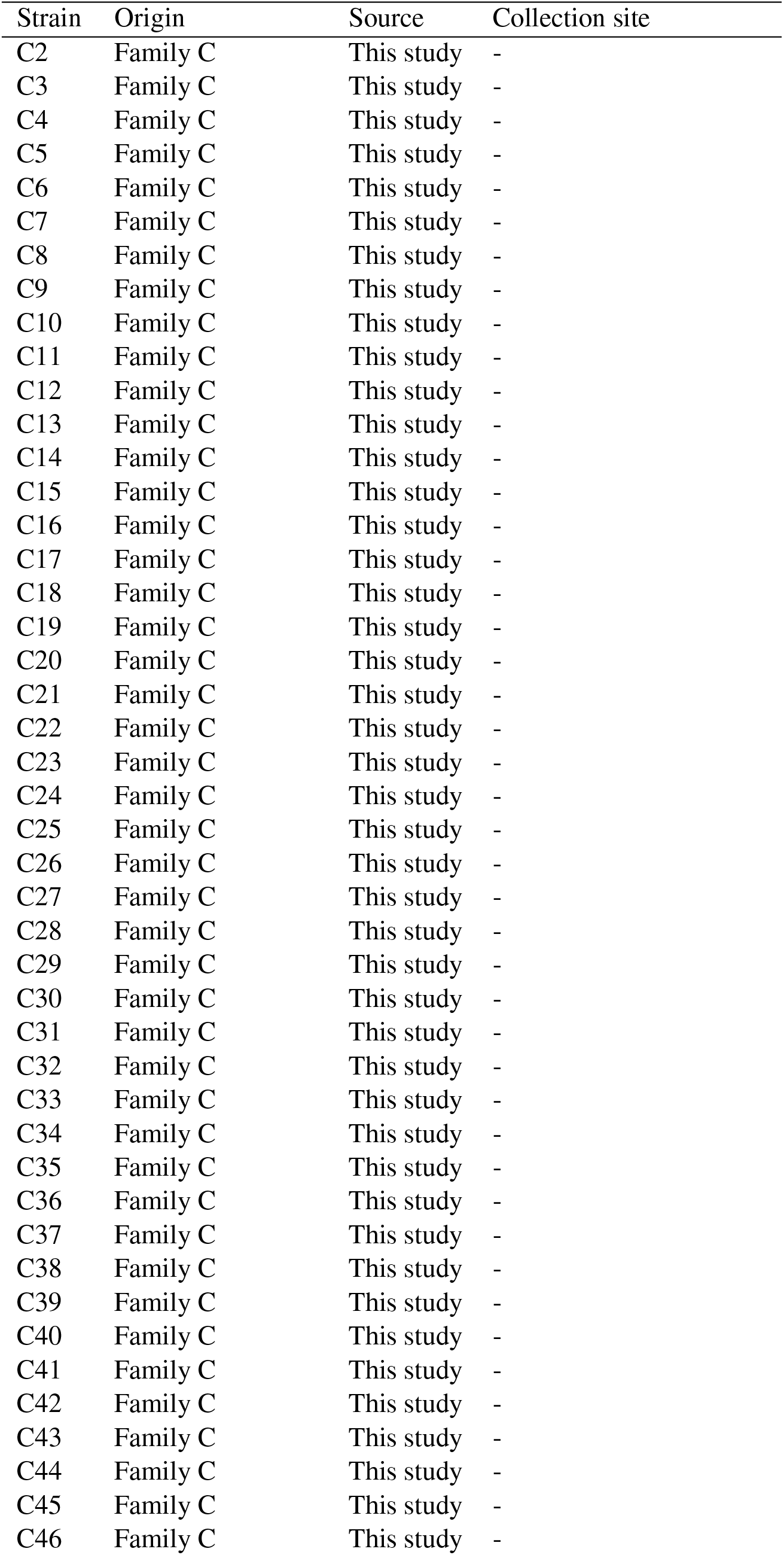
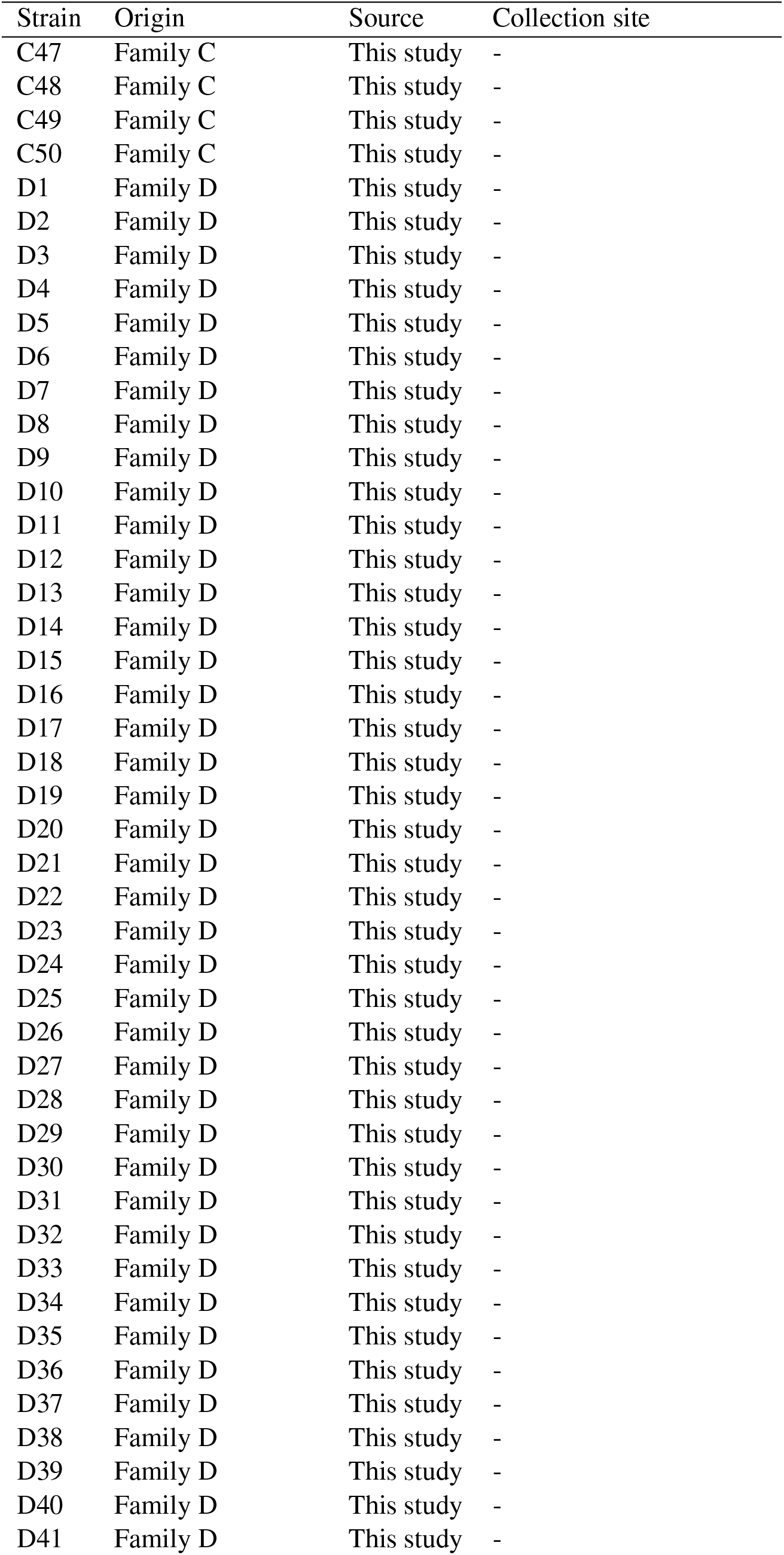
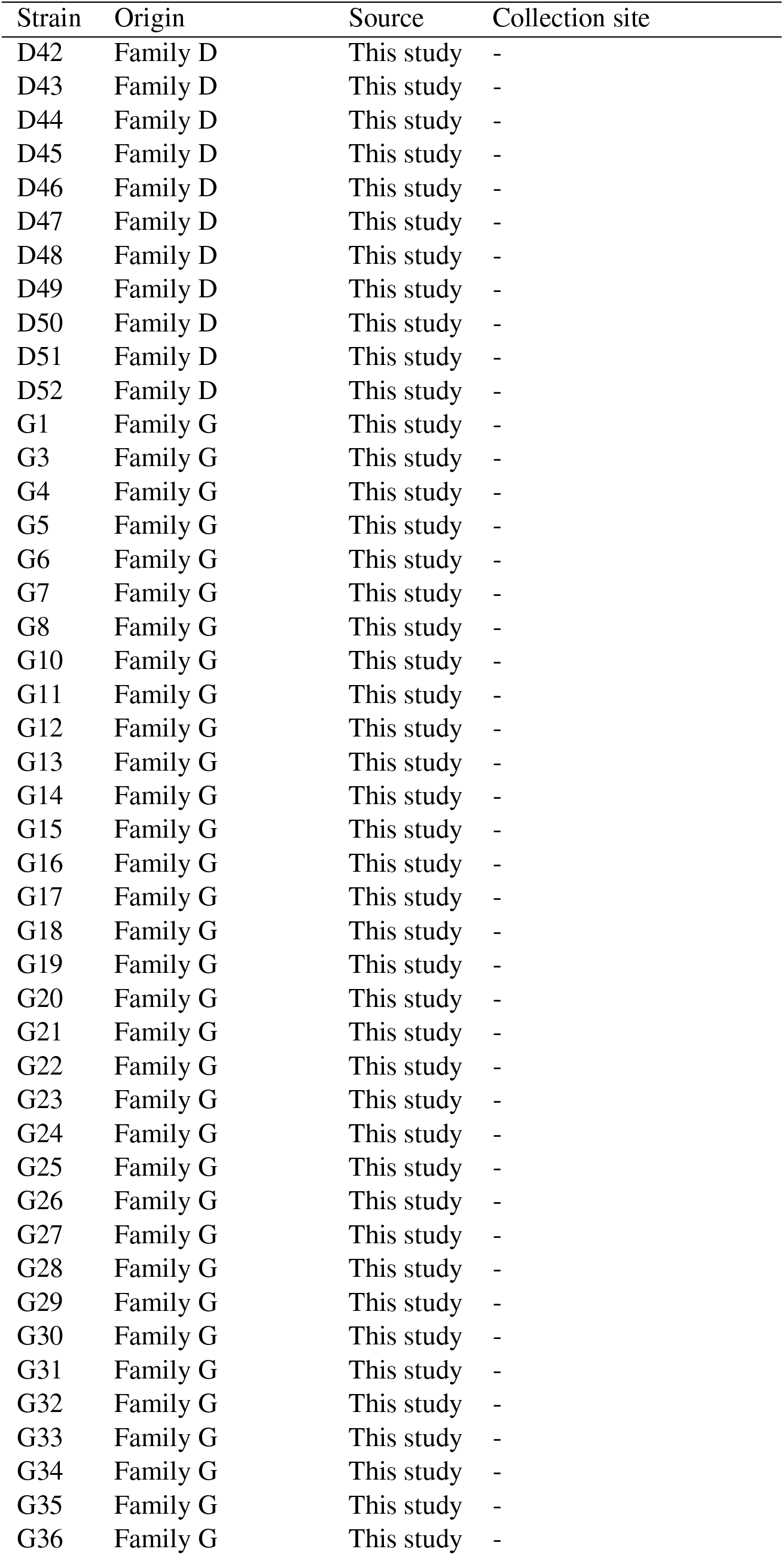
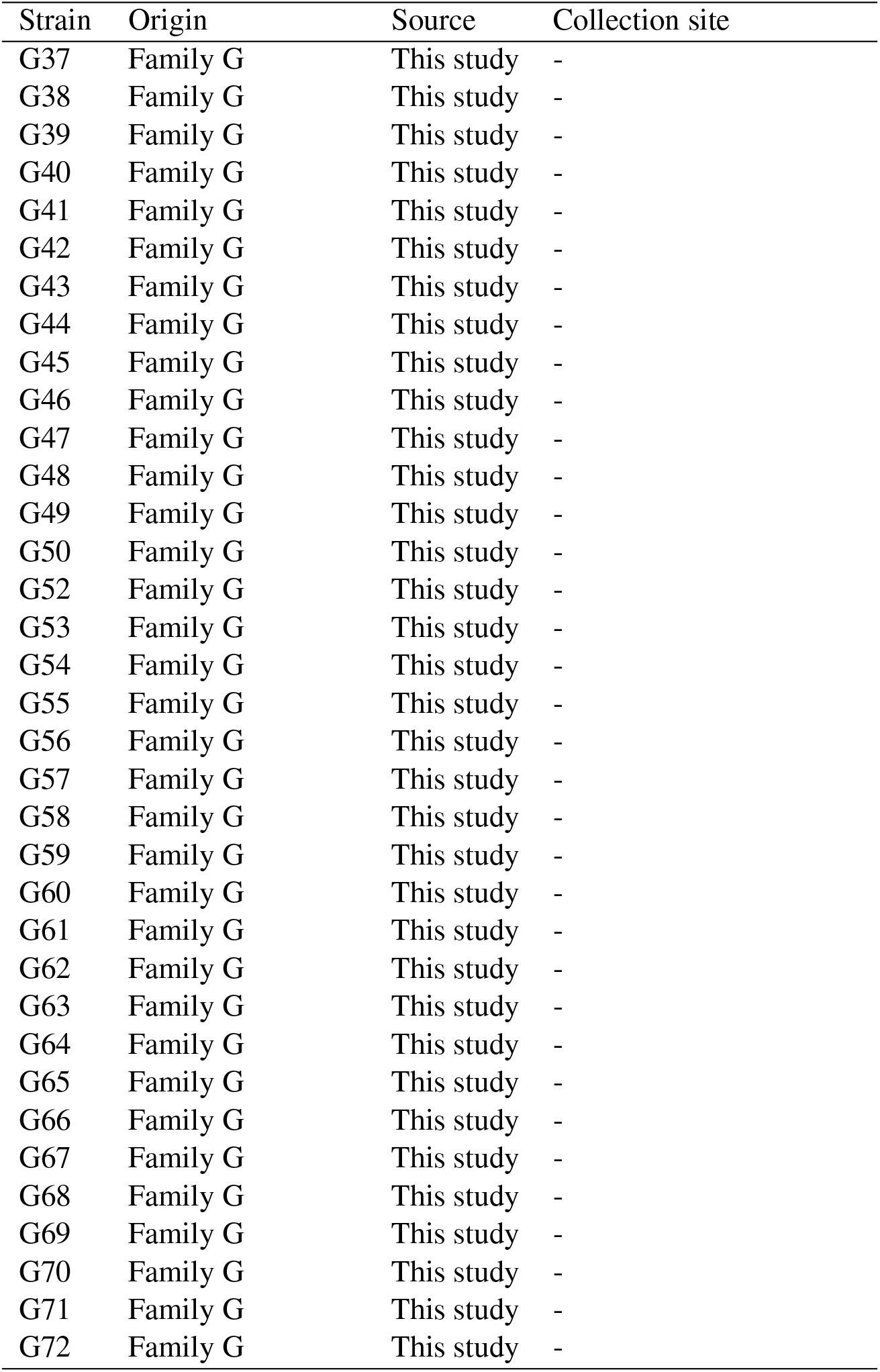
List of strains. Column origin indicates wheter strain was sampled from a natural population or if it was from a family obtained by crossing two natural strains. Column source indicates which strains were obtained from the Fungal Genetics Stock Center (FGSC) and which were generated in this study. LA = Louisiana, USA. FL = Florida, USA. Strains 10948 and 10886 are parents of family A, 10932 and 1165 are parents of family B, 4498 and 8816 are parents of family C, 3223 and 8845 are parents of family D, and 10904 and 851 are parents of family G.

## Notes

### Competing Interest Statement

The authors have declared no competing interest.

### Summary of Updates

We have revised the manuscript following reviewers comments. Biggest changes are that we know use the evolvability statistics of Hansen & Hoyle to evaluate trait autonomy, and how easy it is to evolve along the simulated selection gradients. The simulation part of the manuscript has been extensively revised and figures relating to these results have been redrawn. A better explanation of the biological interpretation of selection and gradient and selection differentials has been added. Furthermore, numerous fixes and clarifications have been done.

